# Molecular Features Underlying Shp1/Shp2 Discrimination by Immune Checkpoint Receptors

**DOI:** 10.1101/2021.06.23.449580

**Authors:** Xiaozheng Xu, Takeya Masubuchi, Yunlong Zhao, Enfu Hui

## Abstract

Numerous inhibitory immunoreceptors operate by recruiting phosphatase effectors Shp1 and Shp2 through conserved motifs ITIM and ITSM. Despite the similarity, these receptors exhibit distinct effector binding specificities, as exemplified by PD-1 and BTLA, which preferentially recruit Shp2 and Shp1 respectively. The molecular basis of Shp1/Shp2 discrimination is unclear. Here, we provide evidence that optimal PD-1 and BTLA binding to both Shp1 and Shp2 occurs via a bivalent, parallel mode that involves both SH2 domains of Shp1/Shp2. Moreover, PD-1 mainly uses its ITSM to discriminate Shp2 from Shp1 via their C-terminal SH2 domains. Supportive of this model, swapping the Shp1-cSH2 with Shp2-cSH2 enabled PD-1:Shp1 association in T cells. In contrast, BTLA primarily utilizes its ITIM to discriminate Shp1 from Shp2 via their N-terminal SH2 domains. Substitution of glycine at pY+1 position of the PD-1-ITIM with alanine, a residue conserved in several Shp1-recruiting receptors, was sufficient to induce PD-1:Shp1 interaction in T cells. Finally, mutagenesis screening shows that Shp1 recruitment exhibits a bell-shaped dependence on the side chain volume of the pY+1 residue of ITIM. Collectively, we provide a molecular interpretation of the Shp1/Shp2-binding specificities of PD-1 and BTLA, with general implications for the mechanism of effector discrimination by inhibitory receptors.

## Main

Signal transduction critically depends on phosphorylation that occurs both on cell surface receptors and on their downstream effector molecules. Many signal-transducing proteins contain Src-Homology-2 (SH2) domains, a structural fold that binds to phosphotyrosine (pY) motifs^1,2^. Thus, tyrosine phosphorylation can induce intermolecular interactions, when pY and SH2 exist in different proteins, or intramolecular interactions, when pY and SH2 exist in the same protein. As such, tyrosine phosphorylation acts as a switch for both protein-protein interactions and protein conformational changes that dictate their functional activities.

In cells, two sets of enzymes reciprocally control the level of tyrosine phosphorylation: protein tyrosine kinases (PTKs) which catalyze the phosphorylation of tyrosine residues, and protein tyrosine phosphatases (PTPases) that catalyze the removal of phosphate groups from pY residues^3-6^. As a prominent example, the T cell antigen receptor (TCR) complex and the costimulatory receptor CD28 both contain phosphorylatable tyrosines in their intracellular domains (ICDs). Phosphorylation of TCR and CD28, catalyzed by the membrane-targeted PTK Lck^7-9^, creates docking sites for SH2-containing, cytoplasmic kinases and adaptor proteins that relay and amplify the receptor signal through well-orchestrated phosphorylation cascades^10,11^.

The phosphorylation-dependent TCR and CD28 signaling is opposed by inhibitory receptors such as PD-1 and BTLA. These receptors recruit cytoplasmic PTPases^12-15^ upon ligation to their respective ligands, PD-L1/2^16-18^ and HVEM^19-21^, to dephosphorylate TCR and CD28 signaling components^22-24^. Both PD-1 and BTLA contain a single ITIM (immunoreceptor tyrosine-based inhibition motif; S/I/V/LxYxxI/V/L) and a single ITSM (immunoreceptor tyrosine-based switch motif; TxYxxV/I), conserved motifs found in many immunoreceptors^25-28^. Upon tyrosine phosphorylation, ITIM and ITSM serve as docking sites for tandem SH2 domains (tSH2) contained in Shp1 and Shp2, structurally similar, and yet functionally distinct cytoplasmic PTPases^29^. The PTPase domain of Shp1, but not that of Shp2, was able to dephosphorylate Zap70 and Syk^30^. Moreover, Shp1-recruiting receptors more potently suppress TCR and CD28 phosphorylation than do Shp2-recruiting receptors^24^.

Despite the large number of ITIM/ITSM-containing receptors in the human genome, little is known regarding their molecular mechanisms. There is sparse information about their specificities for Shp1 or Shp2. Whether they signal through Shp1, Shp2, or both, and to what extent they prefer one over the other, have not been well established. Indeed, growing evidence suggests that ITIM/ITSM-containing receptors exhibit distinct specificities for Shp1 and Shp2 recruitment. PD-1 strongly recruits Shp2, but not Shp1, in both T cells and B cells^14,23^. In contrast, BTLA prefers Shp1 over Shp2^24,31,32^. The mechanism underlying the distinct PTPase-binding specificities of PD-1 and BTLA remains obscure. This issue has been confounded by the multiple possible binding modes between a dual-tyrosine receptor and a dual-SH2 protein, and the reported abilities of the Shp1/Shp2 to dephosphorylate their docking sites within the receptor^22,23,33^.

In this study, we dissected the molecular mechanisms by which PD-1 and BTLA discriminate between Shp1 and Shp2. We measured the affinities of all potential pY:SH2 interactions involved in PD-1 and BTLA recruitment of Shp1 and Shp2 using surface plasmon resonance (SPR), and identified the optimal binding geometries of both Shp1 and Shp2. We then measured the recruitment of Shp1 and Shp2 to PD-1 microclusters in intact T cells, using both wild-type (WT) and domain-swapped mutants. These experiments have allowed us to identify key molecular features in PD-1 and BTLA that lead to their specificity dichotomy. Our finding has implications to the specificities of the rapidly growing list of immune checkpoint receptors.

## Results

### PD-1 recruits Shp2, but not Shp1, whereas BTLA prefers to recruit Shp1

Previous studies suggest that tyrosine-phosphorylated PD-1 recruits Shp2, but not Shp1, to suppress T cell activation^23,34^. To begin investigating the molecular basis of this specificity, we utilized an antigen presenting cell (APC) – T cell co-culture assay incorporating the PD-L1:PD-1 pathway. In this assay, PD-1-mGFP transduced Jurkat T cells were stimulated with superantigen (SEE)-loaded, PD-L1 transduced Raji B cells (APCs). After lysing the cell conjugates at desired time points, we immunoprecipitated (IP) PD-1-mGFP from the cell lysates and probed pY and co-precipitated Shp1 or Shp2 using immunoblots (IB). PD-1 became tyrosine phosphorylated, and recruited Shp2 but not Shp1 in a time-dependent fashion (**Fig. 1a**), consistent with recent studies^23,24^. By contrast, in a parallel co-culture system containing HVEM transduced Raji cells and BTLA-mGFP transduced Jurkat cells, BTLA recruited both Shp1 and Shp2, with a clear preference for Shp1 (**Fig. 1b**), consistent with recent studies^24,31,32^.

**Fig. 1:**
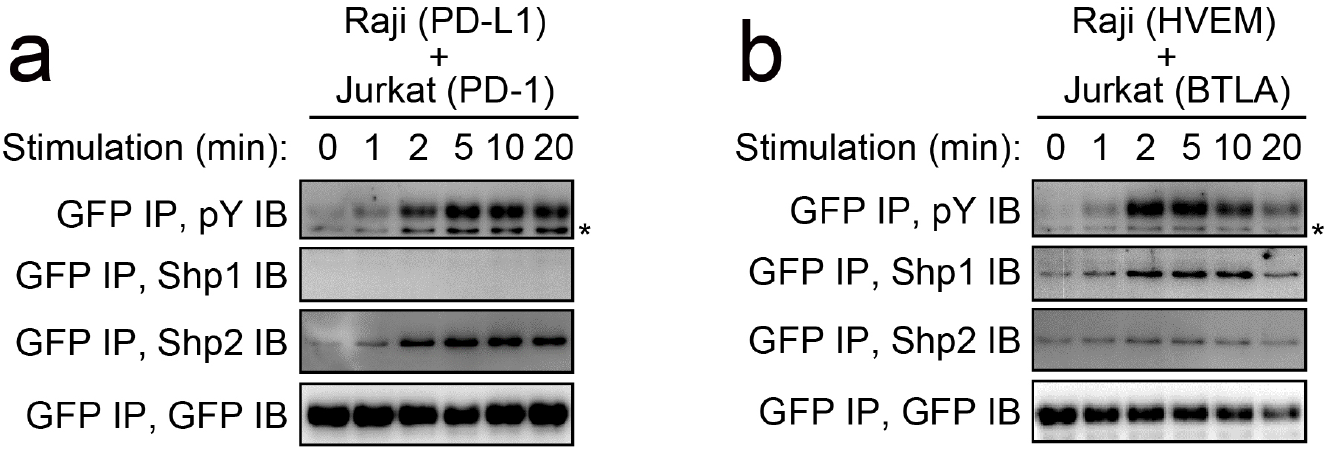
PD-1 recruits Shp2 but not Shp1, whereas BTLA prefers to recruit Shp1. Representative immunoblots (IBs) showing the levels of bound Shp1 and Shp2 in PD-1-mGFP (**a**) or BTLA-mGFP (**b**) pulled down by GFP IP from indicated cell lysates, with the duration of stimulation prior to lysis indicated (see Methods). IBs of GFP and pY of the same samples were shown to indicate PD-1 or BTLA input and their degrees of phosphorylation.

### Both ITIM and ITSM contribute to the ability of PD-1 to recruit Shp2

We next attempted to clarify the relative contributions of ITIM and ITSM in mediating Shp2 recruitment, by examining how mutations of these motifs affect Shp2 binding and PD-1 function. We generated Jurkat cells expressing similar levels of mGFP-tagged PD-1^WT^, PD-1^FY^, in which the ITIM Y223 was mutated to phenylalanine, PD-1^YF^, in which the ITSM Y248 was mutated to phenylalanine, or PD-1^FF^, in which both tyrosines were mutated (**Extended Data Fig. 1a**). Upon stimulation with PD-L1-transduced Raji cells, we detected Shp2 but not Shp1 in the PD-1^WT^ IP (GFP IP), and as expected, Shp2 was undetectable in PD-1^FF^ IP samples. The ITSM mutant PD-1^YF^ also failed to recruit Shp2, whereas the ITIM mutant PD-1^FY^ recruited Shp2, but significantly less than PD-1^WT^ (**Fig. 2a**). We confirmed these observations by visualizing PD-1:Shp2 interaction in intact T cells. We plated the foregoing Jurkat cells on a supported lipid bilayer (SLB) containing anti-CD3ε (for TCR stimulation) and recombinant PD-L1 ectodomain (PD-L1^ECD^, for PD-1 stimulation). Total internal reflection microscopy (TIRF-M) in the GFP channel revealed PD-1 microclusters in all four cell-types (**Fig. 2b**). Immunostaining of Shp2 showed strong enrichment of Shp2 to PD-1^WT^ microclusters. Shp2 recruitment was slightly weaker for PD-1^FY^, but statistically significant (p=0.0306), and Shp2 recruitment was almost completely abrogated in PD-1^YF^, similar to the negative control PD-1^FF^ (**Fig. 2b**). These data are in general agreement with previous reports that ITSM is the dominant docking site for Shp2^12,14,23,35,36^. However, our result showed that PD-1^WT^ recruited more Shp2 than did PD-1^FY^, suggesting that optimal Shp2 recruitment does require ITIM. The more obvious defect of PD-1^FY^ in the co-IP assays might be due to the disruption of weak interactions by the non-equilibrium wash steps.

**Fig. 2:**
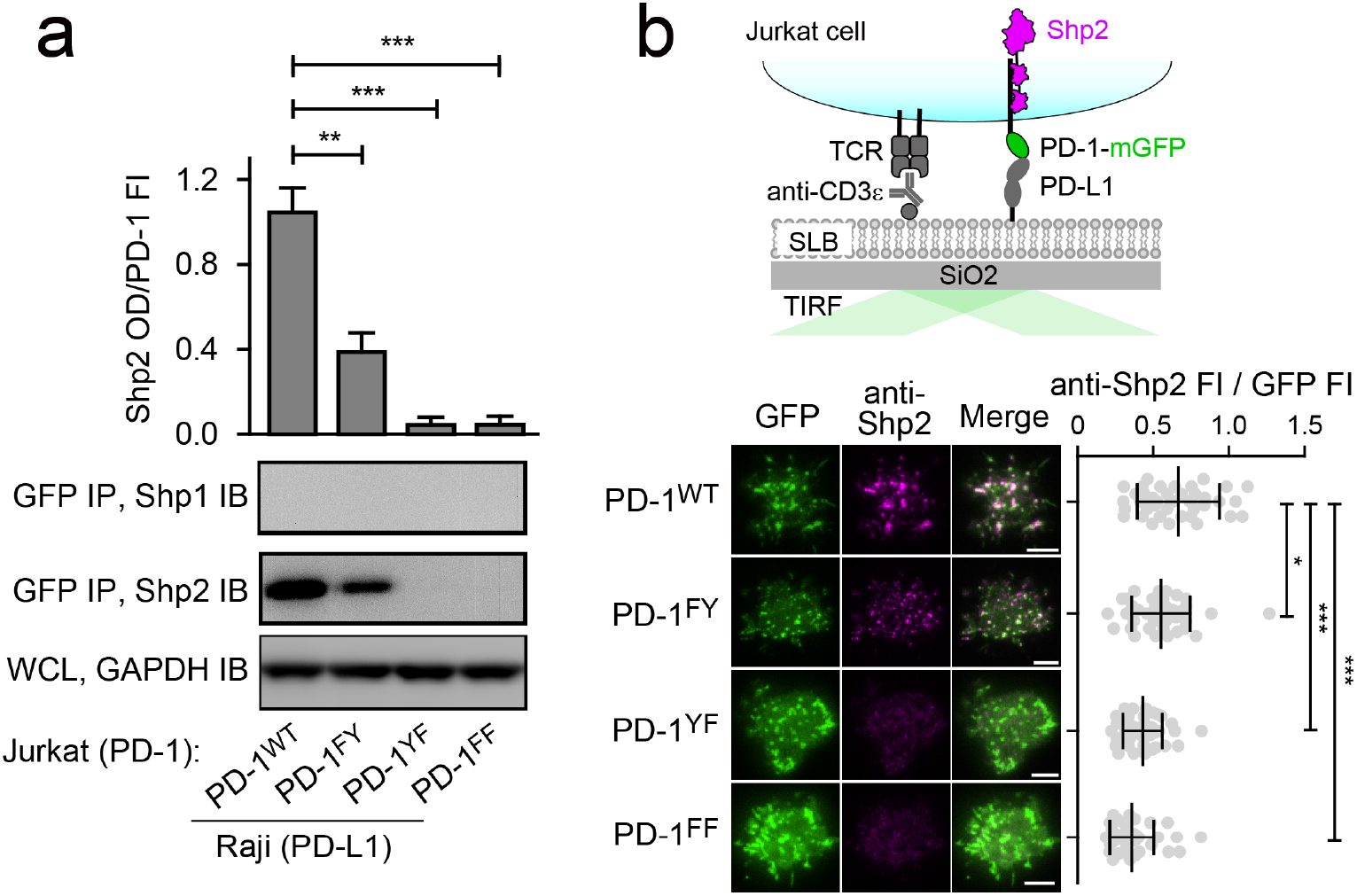
Both ITIM and ITSM contribute to the ability of PD-1 to recruit Shp2. **a**, Representative IBs showing the levels of Shp1 and Shp2 bound to mGFP-tagged PD-1 variants pulled down from the indicated co-culture lysates via GFP IP. GAPDH IB of the whole cell lysates (WCL) served as a loading control (see Methods). Bar graphs on top summarize Shp2 optical density (OD) normalized to the fluorescence intensity (FI) of each PD-1 variant, based on flow cytometry data in Extended Data Fig. 1a. Error bars are s.d. from three independent experiments. **b**, Upper, a cartoon depicting a PD-1-mGFP-expressing Jurkat cell in contact with an SLB containing anti-CD3ε and PD-L1^ECD^. Lower left, representative TIRF images of both PD-1 (GFP) and endogenous Shp2 (stained with anti-Shp2) in an SLB-associated Jurkat expressing indicated PD-1 variants. Lower right, dot plots summarizing anti-Shp2 FI normalized to GFP FI of 40 Jurkat cells under each condition (see Methods); Error bars: s.d.. Scale bars: 5 µm. *P < 0.05; **P < 0.01; ***P < 0.001; ns, not significant; Student’s t-test.

### PD-1-ITSM strongly prefers Shp2-cSH2 over Shp1-cSH2

We next investigated the molecular mechanism by which PD-1 recruits Shp2 but not Shp1 in T cells. Shp1 and Shp2 both contain two SH2 domains in tandem, the N-terminal SH2 (nSH2) and the C-terminal SH2 (cSH2). Our co-IP data indicated that Shp2 interacts with PD-1 in a bivalent fashion involving both SH2 domains. The bivalent interaction can potentially occur either in a parallel fashion in which nSH2 binds to ITIM and cSH2 binds to ITSM, or in an antiparallel fashion in which nSH2 and cSH2 bind to ITSM and ITIM respectively. To determine the most favorable binding orientation, we next measured the affinities for all the possible pY:SH2 interactions implicated in PD-1:Shp1 and PD-1:Shp2 interactions. We purified pre-phosphorylated ICDs of PD-1 mutants that contained only one tyrosine within either ITIM (PD-1^YF^) or ITSM (PD-1^FY^). We then used SPR to measure their binding affinities to purified Shp1-nSH2, Shp1-cSH2, Shp2-nSH2, and Shp2-cSH2 (**Fig. 3a**), and summarized the dissociation constants (*K*_d_) in **Supplementary Table 1**.

**Fig. 3:**
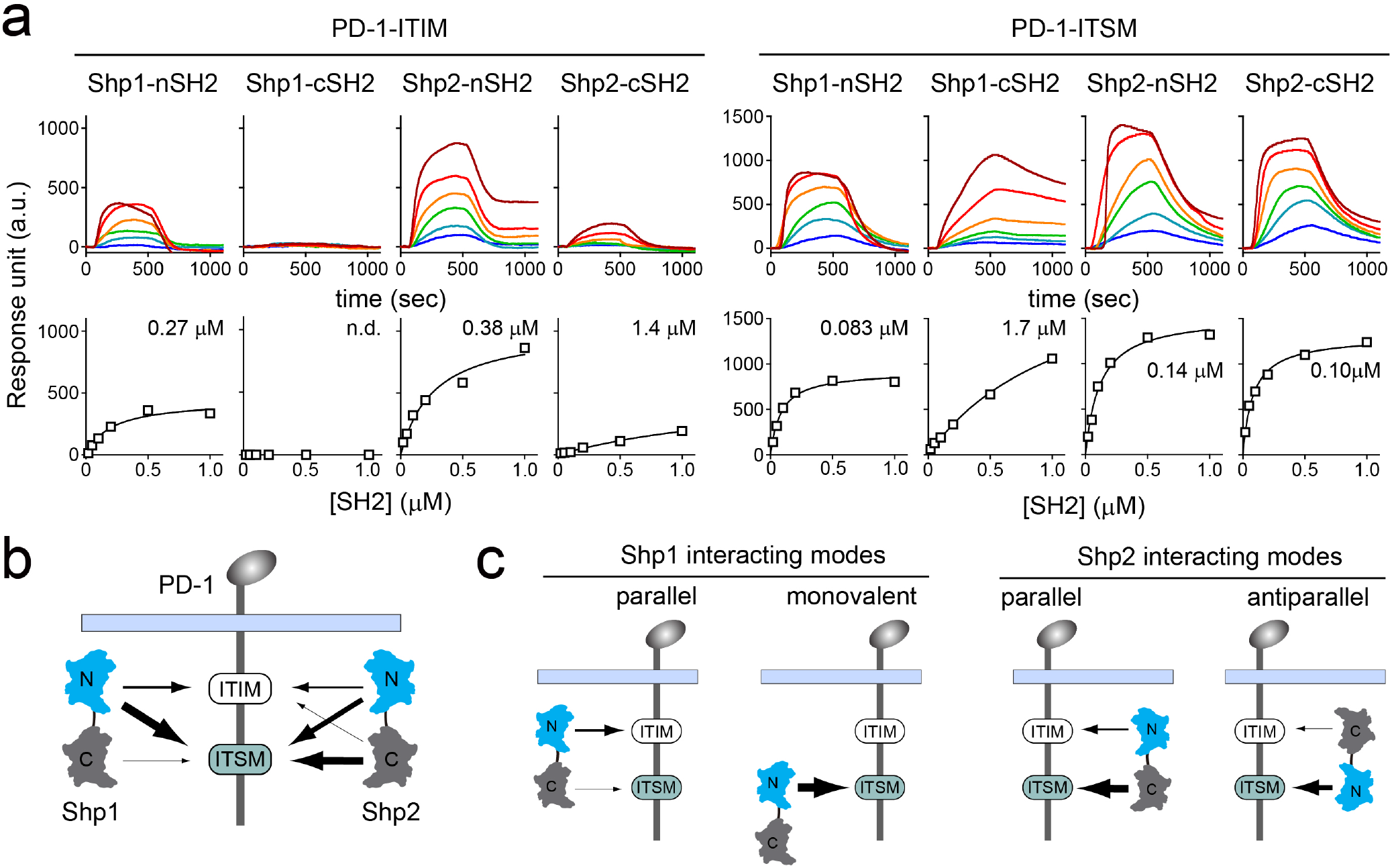
SPR measurements of binding between individual SH2 of Shp1 or Shp2 and ITIM or ITSM of PD-1. **a**, SPR sensorgrams (top) and the derived equilibrium binding curves (bottom) showing the interactions of indicated SH2 and phosphorylated PD-1-ITIM (PD-1 YF) or PD-1-ITSM (PD-1 FY) immobilized onto Ni sensor chips. Individual SH2 proteins were injected at 20, 50, 100, 200, 500, 1000 nM. Shown are representative of three independent experiments. The calculated *K*_d_ values are indicated in the binding curves. **b**, A cartoon depicting relative binding affinities of Shp1/Shp2 individual SH2 to PD-1-ITIM/ITSM, with the thickness of arrows matching the relative affinities calculated from the SPR data. **c**, Possible interacting modes of Shp1/Shp2-tSH2 with PD-1.

To summarize the SPR data, we diagramed all the detectable SH2:pY interactions in the context of tandem SH2 and PD-1^WT^, with relative affinities depicted by arrow thickness. PD-1-ITSM is a better docking site than is PD-1-ITIM for each of the four SH2 tested (**Fig. 3b**). A careful inspection of the data also revealed specific information, as detailed below.

For PD-1:Shp2 interactions, Shp2-nSH2 weakly preferred PD-1-ITSM (*K*_d_ = 0.14 μM) over PD-1-ITIM (*K*_d_ = 0.38 μM), and Shp2-cSH2 strongly preferred PD-1-ITSM (*K*_d_ = 0.10 μM) over PD-1-ITIM (*K*_d_ = 1.4 μM) (**Fig. 3a**,**b**; **Supplementary Table 1**). Thus, the parallel mode PD-1:Shp2 complex would be more energetically favorable than the antiparallel mode (**Fig. 3c**, right; **Supplementary Tables 2** and **3**), consistent with a recent report^37^.

For PD-1:Shp1 interactions, Shp1-nSH2 preferred PD-1-ITSM (*K*_d_ = 0.083 μM) over PD-1-ITIM (*K*_d_ = 0.27 μM). Interestingly, Shp1-cSH2 appeared to be defective in PD-1 binding, exhibiting a rather weak affinity to PD-1-ITSM (*K*_d_ = 1.7 μM), and no detectable binding to PD-1-ITIM (**Fig. 3a,b**; **Supplementary Table 1**). The inability of Shp1-cSH2 to bind PD-1-ITIM ruled out the anti-parallel mode of PD-1:Shp1 interactions, but indicated the possibility of a monovalent mode in which Shp1-nSH2 interacts with PD-1-ITSM (**Fig. 3c**, left). However, free energy calculations suggested that the parallel mode, which involves two SH2, is energetically favorable over the monovalent mode (**Supplementary Table 2** and **3**).

To further examine the dominant mode of PD-1:Shp1 interactions, we employed a single molecule assay to determine whether PD-1:Shp1 interactions require only ITSM, as would be expected for the monovalent mode, or both ITIM and ITSM, as would be expected for the bivalent parallel mode. We sparsely attached monomeric, fluorescently-labeled, pre-phosphorylated and biotinylated PD-1^WT^, PD-1^FY^, or PD-1^YF^ (**Extended Data Fig. 2a,b**) to a biotin polyethylene glycol (PEG)-coated coverslip via streptavidin. TIRF-M resolved individual PD-1 monomers as discrete spots, which underwent photobleaching in single steps (**Extended Data Fig. 2c,d**). After the addition of JF646-labeled tSH2 of either Shp1 or Shp2, we visualized PD-1:tSH2 interaction at the coverslip (**Extended Data Fig. 2e**, left). Recruitment of Shp2-tSH2 to PD-1 led to the appearance of JF646 signal that colocalized with PD-1 molecules (**Extended Data Fig. 2e**, upper right). Each Shp2-tSH2 spot typically persisted for several seconds, then disappeared due to dissociation from PD-1, leading to a step-like time course (**Extended Data Fig. 2e**, lower right). As expected, Shp1-tSH2 displayed a lower degree of PD-1 occupancy (**Extended Data Fig. 2f**). Moreover, mutation of either ITIM (PD-1^FY^) or ITSM (PD-1^YF^) strongly reduced the PD-1 occupancy for both Shp1-tSH2 and Shp2-tSH2, as compared to the WT control (PD-1^WT^) (**Extended Data Fig. 2f**), further supporting that both Shp1-tSH2 and Shp2-tSH2 binds to PD-1^WT^ in a bivalent fashion involving both ITIM and ITSM.

Collectively, data presented in this section demonstrated that both Shp1 and Shp2 interact with PD-1 primarily via the bivalent parallel geometry. However, the PD-1:Shp1 interaction is much less stable due to the very weak affinity between Shp1-cSH2 and PD-1-ITSM.

### Replacement of Shp1-cSH2 by Shp2-cSH2 is sufficient to induce PD-1:Shp1 association in T cells

The SPR data (**Fig. 3b**; **Supplementary Table 1**) indicated that the Shp1-cSH2 barely interacts with PD-1-ITSM, whereas Shp2-cSH2 displayed a 17-fold higher affinity to the ITSM of PD-1. Thus, we next determined if swapping the cSH2 of Shp1 with that of Shp2 could induce PD-1:Shp1 binding in T cells. We sought to image the recruitment of domain-swapped chimeric mutants of Shp1 to PD-1 microclusters. To avoid competition from endogenous Shp1 and Shp2, we generated Shp1/Shp2 double knockout (DKO) Jurkat cells, and co-transduced PD-1-mGFP with mCherry-tagged Shp1^WT^ or Shp1 mutant with one or both of its SH2 domains replaced by those of Shp2 (**Fig. 4a**). Having confirmed that these cells expressed similar levels of PD-1-mGFP and mCherry-tagged Shp1 variants (**Extended Data Fig. 1b**), we stimulated each type of Jurkat cells with a SLB containing anti-CD3ε and PD-L1^ECD^.

**Fig. 4:**
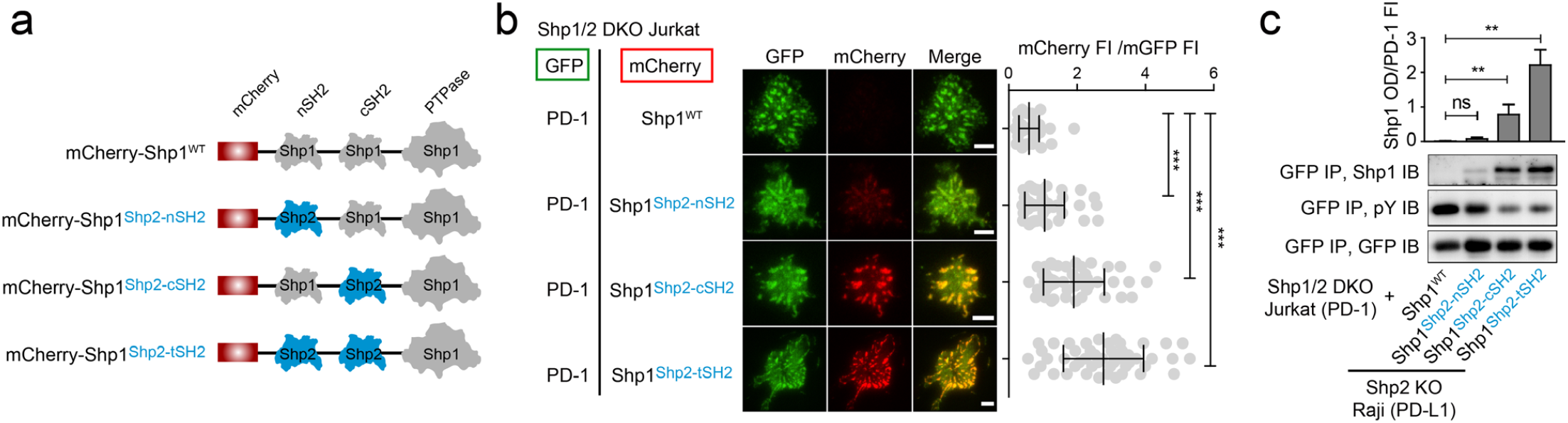
Swapping the cSH2 of Shp1 with that of Shp2 induced PD-1:Shp1 association in T cells. **a**, Diagram showing the design of mCherry-tagged, SH2-swapped Shp1 variants, with one or both of its SH2 replaced with that of Shp2. **b**, Left, representative TIRF images of PD-1 (GFP) and Shp1 variants (mCherry) in SLB-associated Shp1/Shp2 DKO Jurkat expressing PD-1-mGFP and mCherry-Shp1 variants. Right, dot plots summarizing mCherry FI normalized to GFP FI of 40 Jurkat cells under each condition. Error bars: s.d.. Scale bars: 5 µm. **c**, Representative IBs of mCherry-Shp1 variants co-precipitated with PD-1-mGFP from indicated cell lysates. IBs of GFP indicate PD-1 input. IBs of pY indicate PD-1 phosphorylation. Bar graphs summarize OD of Shp1 variants normalized to the FI of PD-1, based on flow cytometry data in Extended Data Fig.1b. Error bars are s.d. from three independent experiments. *P < 0.05; **P < 0.01; ns, not significant; Student’s t-test.

TIRF-M showed that upon cell-bilayer contact, PD-1 formed microclusters that recruited little to no Shp1^WT^, as manifested by nearly undetectable mCherry signal in the GFP foci (**Fig. 4b**, top row), consistent with previous reports^23,24^. Swapping the nSH2 of Shp1 with that of Shp2 (Shp1^Shp2-nSH2^) increased the mCherry signal in PD-1 microclusters, but only to a minor degree (**Fig. 4b**, second row). In contrast, swapping the cSH2 of Shp1 with that of Shp2 (Shp1^Shp2-cSH2^) led to a marked increase in mCherry signal in the PD-1 microclusters (**Fig. 4b**, third row), to a comparable extent as Shp1^Shp2-tSH2^, in which both SH2 of Shp1 were replaced with those of Shp2 (**Fig. 4b**, bottom row).

We confirmed the TIRF results using a co-IP assay. After stimulation of the foregoing Jurkat cells (**Fig. 4b**) with PD-L1-transduced Raji cells, we pulled down PD-1-mGFP and examined the co-precipitated Shp1 variants using IB. We detected no signal of Shp1^WT^, weak signal of Shp1^Shp2-nSH2^, and strong signal of Shp1^Shp2-cSH2^ and Shp1^Shp2-tSH2^ (**Fig. 4c**, GFP IP, Shp1 IB). Notably, PD-1 phosphorylation was inversely correlated with the recruitment of the Shp1 variants (**Fig. 4c**, GFP IP, pY IB), supporting the notion that PD-1 is also a substrate for its bound PTPases^22,23,33^.

Collectively, data reported in this section demonstrated that cSH2 is the major determinant underlying PD-1’s strong preference for Shp2 over Shp1.

### ITIM and ITSM are both required for BTLA to recruit Shp1/Shp2

Having established deficient ITSM:cSH2 interactions as the primary basis for the reduced stability of PD-1:Shp1 binding, we next turned our attention to receptors that normally recruit Shp1 in T cells, such as BTLA (**Fig. 1b**) to gain further insights into the mechanisms of effector PTPase discrimination by ITSM/ITIM-bearing receptors^13,24,31^. We wished to determine the structural features in BTLA that enabled its Shp1 recruitment in T cells.

The ICD of human BTLA harbors four phosphorylatable tyrosines (Y226, Y243, Y257 and Y282), in which Y257 and Y282 are embedded in ITIM and ITSM respectively. We first asked which tyrosine of BTLA are required for Shp1 and Shp2 recruitment. Analogous to PD-1 assays (**Fig. 2a**), we established Jurkat cell lines expressing either WT or mutant BTLA in which one of the four phosphorylatable tyrosine was replaced by phenylalanine (**Fig. 5a**). We then stimulated these cell lines, in parallel, with HVEM-expressing Raji B cells. Co-IP experiments showed that the mutation of either or both of the non-ITIM/ITSM phosphorylatable tyrosines (Y226F: BTLA^FYYY^; Y243F: BTLA^YFYY^; Y226F & Y243F: BTLA^FFYY^) had little to no effect on the abilities of BTLA to recruit Shp1/Shp2 (**Fig. 5b** and **Extended Data Fig. 3**), demonstrating that these two tyrosines are dispensable for BTLA-mediated recruitment of Shp1/Shp2. In contrast, mutation of either ITIM tyrosine (Y257F: BTLA^YYFY^) or ITSM tyrosine (Y282F: BTLA^YYYF^) abolished the binding of both Shp1 and Shp2 (**Fig. 5b**), consistent with previous studies^13^. Thus, ITIM and ITSM are both necessary for Shp1 and Shp2 recruitment by BTLA. These data also suggest that both SH2 domains of Shp1 and Shp2 are required for their recruitment to BTLA. The more stringent requirement of both ITIM and ITSM of BTLA indicates that its ITIM and ITSM play more balanced roles in mediating Shp1/Shp2 recruitment than those of PD-1.

**Fig. 5:**
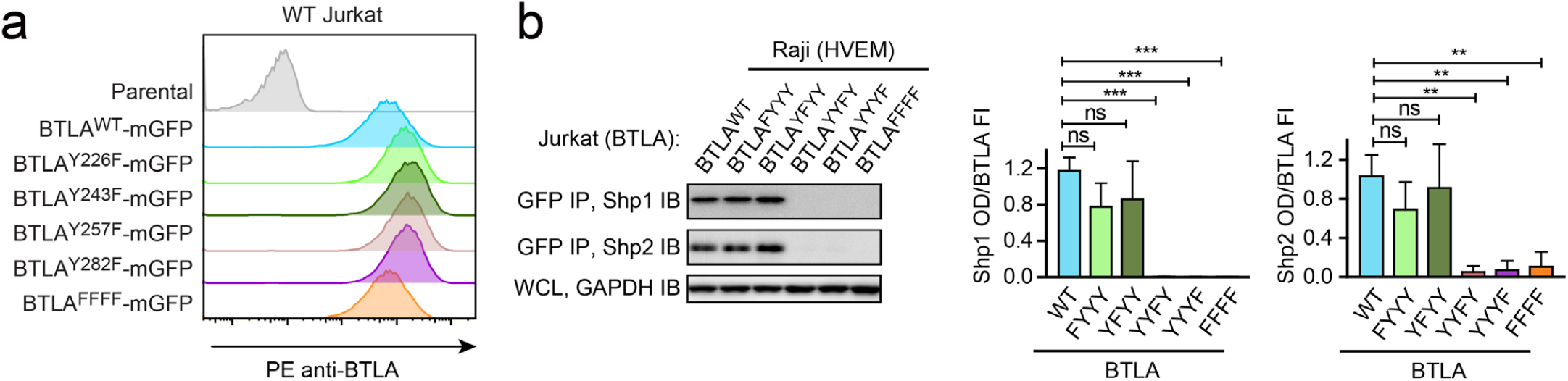
Both ITIM and ITSM are required for BTLA to recruit Shp1/Shp2. **a**, Flow cytometry histograms showing BTLA surface expressions in the indicated Jurkat cells. **b**, Left, representative IBs showing the levels of Shp1 and Shp2 bound to the mGFP-tagged BTLA variants captured by GFP IP. GAPDH IB of the WCL served as a loading control. Right, bar graphs summarizing Shp1 OD and Shp2 OD normalized to the FI of the corresponding BTLA variants, based on flow cytometry data in A. Error bars are s.d. of three independent experiments. **P < 0.01; ***P < 0.001; ns, not significant; Student’s t-test (n = 3).

### Shp1 and Shp2 both interact with BTLA via the bivalent parallel mode

We next wished to determine the most favorable binding geometries for BTLA:Shp1 interactions and BTLA:Shp2 interactions. Analogous to SPR assays with PD-1 (**Fig. 3**), we measured the affinities of individual pY:SH2 interaction implicated in BTLA:Shp1/Shp2 interactions, using sensor chips coated with pre-phosphorylated BTLA triple-tyrosine-mutant FFYF (BTLA-ITIM), which contained a lone tyrosine (Y257) in its ITIM, or pre-phosphorylated BTLA triple-tyrosine-mutant FFFY (BTLA-ITSM), which contained a lone tyrosine (Y282) in its ITSM (**Fig. 6a**). These experiments revealed that for both Shp1 and Shp2, their nSH2 and cSH2 prefer BTLA-ITIM and BTLA-ITSM, respectively (**Fig. 6a,b**; **Supplementary Table 1**). Thus, we concluded that the most favorable BTLA:Shp1 and BTLA:Shp2 interactions both occur in a parallel mode (**Fig. 6c**), similar to PD-1:Shp1 and PD-1:Shp2 interactions.

**Fig. 6:**
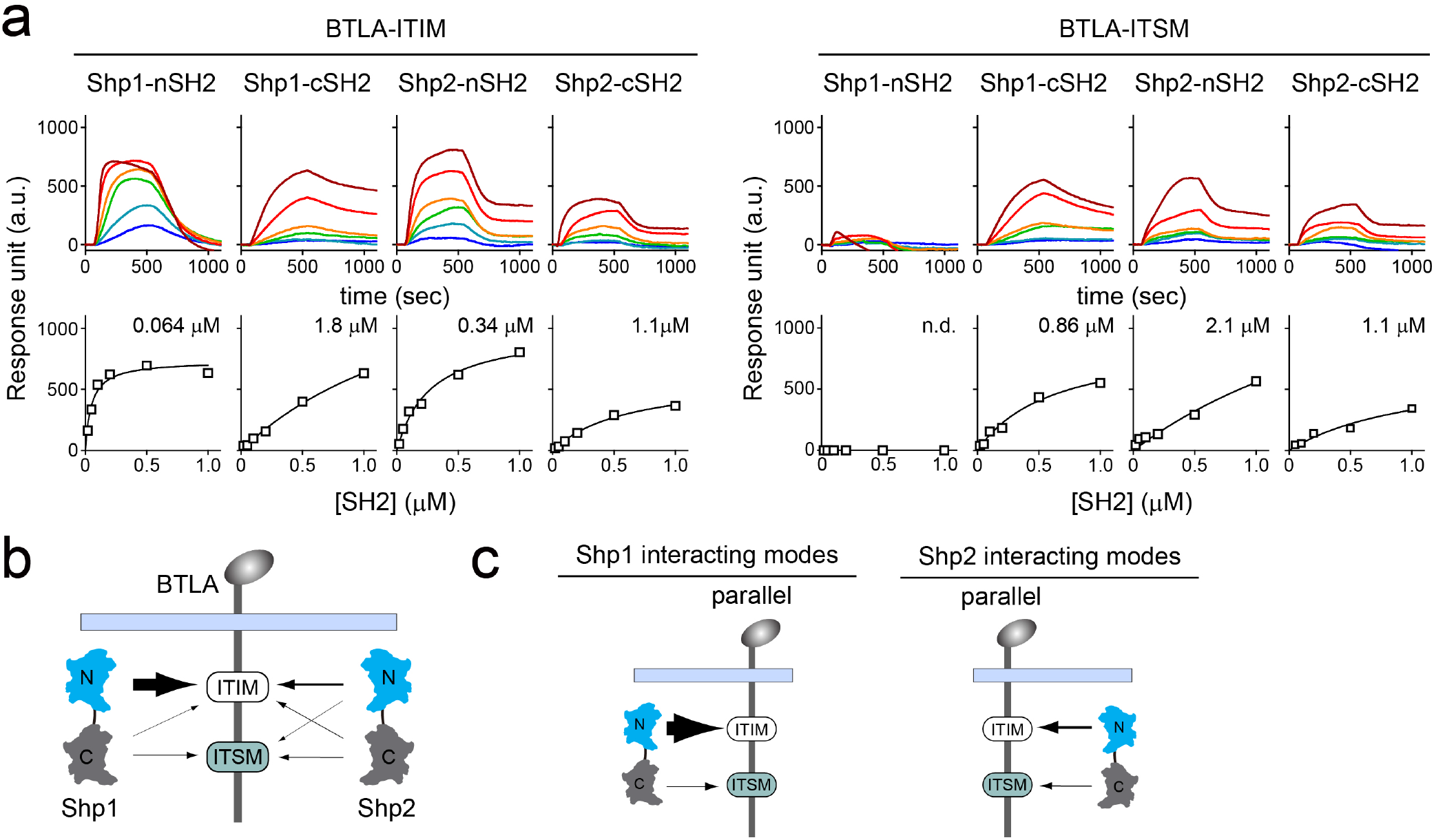
SPR measurements of binding between individual SH2 of Shp1 or Shp2 and ITIM or ITSM of BTLA. **a**, SPR sensorgrams (top) and the derived equilibrium binding curves (bottom) showing the interactions of indicated SH2 and phosphorylated BTLA-ITIM (BTLA FFYF) or BTLA-ITSM (BTLA FFFY) immobilized to Ni sensor chips. Individual SH2 proteins were injected at 20, 50, 100, 200, 500, 1000 nM. Shown are representative of three independent experiments. The calculated *K*_d_ values are indicated in the binding curves. **b**, A cartoon depicting relative binding affinities of Shp1/Shp2 individual SH2 to BTLA-ITIM/ITSM, with the thickness of arrows matching the relative affinities calculated from the SPR data in A. **c**, Possible interacting modes of Shp1/Shp2-tSH2 with BTLA.

### BTLA-ITIM is a high-affinity docking site for Shp1-nSH2

On a closer inspection of the SPR data, we found that the relative contribution of ITIM and ITSM in BTLA is opposite to that in PD-1. While the ITSM is the major SH2 docking site in PD-1, the ITIM appeared to be the major SH2 docking site in BTLA. This is particularly striking in the case of BTLA:Shp1 interaction: Shp1-nSH2 exhibited an impressive affinity to BTLA-ITIM (*K*_d_ = 0.064 μM), 13-fold higher than the affinity between Shp1-cSH2 and BTLA-ITSM (*K*_d_ = 0.86 μM) (**Fig. 6a** and **Supplementary Table 1**).

Notably, the affinity of Shp1-nSH2 to BTLA-ITIM (*K*_d_ = 0.064 μM) was also four-fold higher than its affinity to PD-1-ITIM (*K*_d_ = 0.27 μM) (**Fig. 6a** and **Supplementary Table 1**). Thus, even though both BTLA-ITSM and PD-1-ITSM are poor docking sites for Shp1-cSH2, BTLA-ITIM is a much better docking site for Shp1-nSH2 than is PD-1-ITIM. Conceivably, the strong BTLA-ITIM:Shp1-nSH2 interaction may compensate the weak BTLA-ITSM:Shp1-cSH2 interaction, leading to an overall stable BTLA:Shp1 association in T cells.

### Swapping PD-1-ITIM with BTLA-ITIM induced PD-1:Shp1 interaction in T cells

The foregoing data support a hypothesis that the stability of ITIM:SH2 interactions is vital for ITIM/ITSM-bearing receptors to recruit Shp1. To test this experimentally, we assayed whether replacing the “low affinity” ITIM of PD-1 with the ‘high affinity’ ITIM of BTLA could induce PD-1:Shp1 interaction. We transduced Jurkat cells with either mGFP-tagged PD-1^WT^ or PD-1^BTLA-ITIM^, in which we replaced PD-1-ITIM (VD<u>Y</u>GEL) with BTLA-ITIM (IV<u>Y</u>ASL) (**Fig. 7a** and **Extended Data Fig. 1c**). Following stimulation of both types of Jurkat cells using PD-L1-expressing Raji B cells, co-IP assays revealed that PD-1^BTLA-ITIM^, but not PD-1^WT^ recruited Shp1 (**Fig. 7b**). We confirmed this finding using Shp2 KO Jurkat cells, which allowed us to examine PD-1:Shp1 interaction without potential competition from Shp2 (**Fig. 7a,b**). We further verified these findings in intact cells using TIRF-M. In the cell-SLB assays, we observed microclusters of both PD-1^WT^ and PD-1^BTLA-ITIM^, and confirmed that the PD-1^BTLA-ITIM^ microclusters recruited significantly more Shp1 than did the PD-1^WT^ microclusters (**Fig. 7c**). Finally, in a reciprocal set of experiments, we found that replacement of the BTLA-ITIM with the PD-1-ITIM markedly decreased the Shp1 recruitment to BTLA, in both WT Jurkat and Shp2 KO Jurkat cells (**Extended Data Fig. 4**). Together, data presented in this section demonstrated that Shp1 primarily discriminates BTLA from PD-1 based on their ITIMs.

**Fig. 7:**
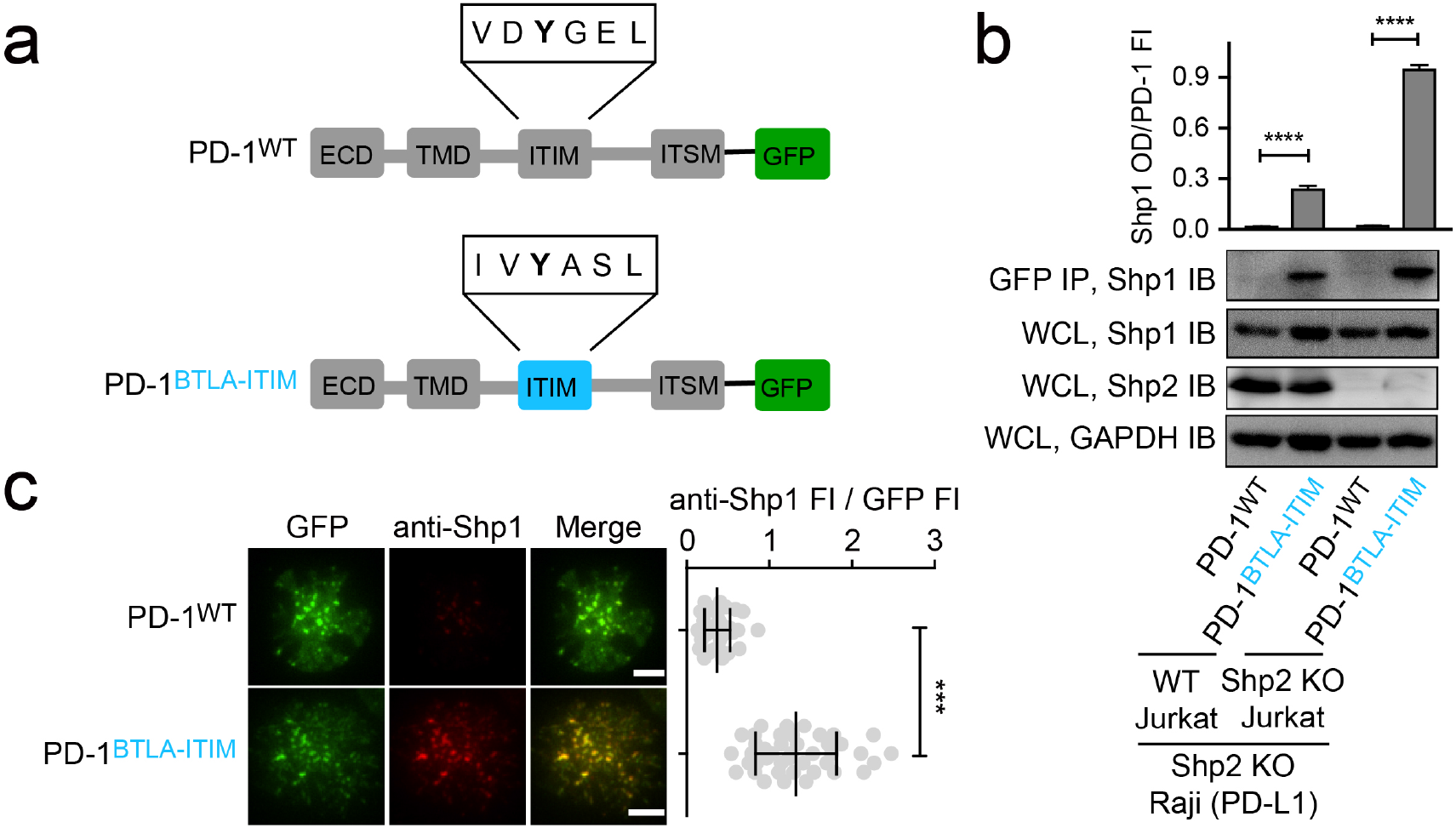
Swapping PD-1-ITIM with BTLA-ITIM induced PD-1:Shp1 interaction in T cells. **a**, Cartoons depicting the domains and motifs of PD-1^WT^-mGFP and PD-1^BTLA-ITIM^-mGFP. **b**, Representative IBs showing the levels of Shp1 bound to mGFP-tagged PD-1 variants pulled down from the indicated co-culture lysates via GFP IP. Shp1 IB and Shp2 IB of the WCL indicate their input. GAPDH IB of the WCL served as a loading control. Bar graphs summarize Shp1 OD normalized to the FI of each PD-1 variant, based on flow cytometry data in Extended Data Fig. 1c. Error bars are s.d. from three independent experiments. **c**, Left, representative TIRF images of both PD-1 (GFP) and endogenous Shp1 (stained with anti-Shp1) in an SLB-associated Shp2 KO Jurkat expressing indicated PD-1 variants. Right, dot plots summarizing anti-Shp1 FI normalized to GFP FI of 40 Jurkat cells under each condition; Error bars: s.d.. Scale bars: 5 μm. ***P < 0.001; ****P < 0.0001; Student’s t-test.

### Replacement of the pY+1 glycine in PD-1-ITIM with alanine was sufficient to induce PD-1:Shp1 association in T cells

We noted that the BTLA-ITIM (IV<u>Y</u>ASL) differ from PD-1-ITIM (VD<u>Y</u>GEL) at four residues flanking pY: V221, D222, G224 and E225 in PD-1-ITIM are replaced by I, V, A and S respectively in BTLA-ITIM (**Fig. 8a**). We wished to determine which replacement contributed the most in inducing Shp1 binding of PD-1^BTLA-ITIM^. We generated Shp2 KO Jurkat cells expressing comparable levels of PD-1^V221I^, PD-1^D222V^, PD-1^G224A^, or PD-1^E225S^, and each harbored a C-terminal mGFP (**Extended Data Fig. 1d**), and asked which mutants were able to recruit Shp1. We also used cells expressing PD-1^WT^-mGFP and cells expressing PD-1^BTLA-ITIM^-mGFP as controls. Upon stimulation with PD-L1-transduced Raji cells, we IP’ed PD-1-mGFP and blotted for Shp1. As expected, Shp1 signal was evident in the precipitates of PD-1^BTLA-ITIM^ but not in PD-1^WT^. Notably, PD-1^G224A^ also recruited the most Shp1 among the single point mutants, despite less than PD-1^BTLA-ITIM^, the positive control (**Fig. 8b**).

**Fig. 8:**
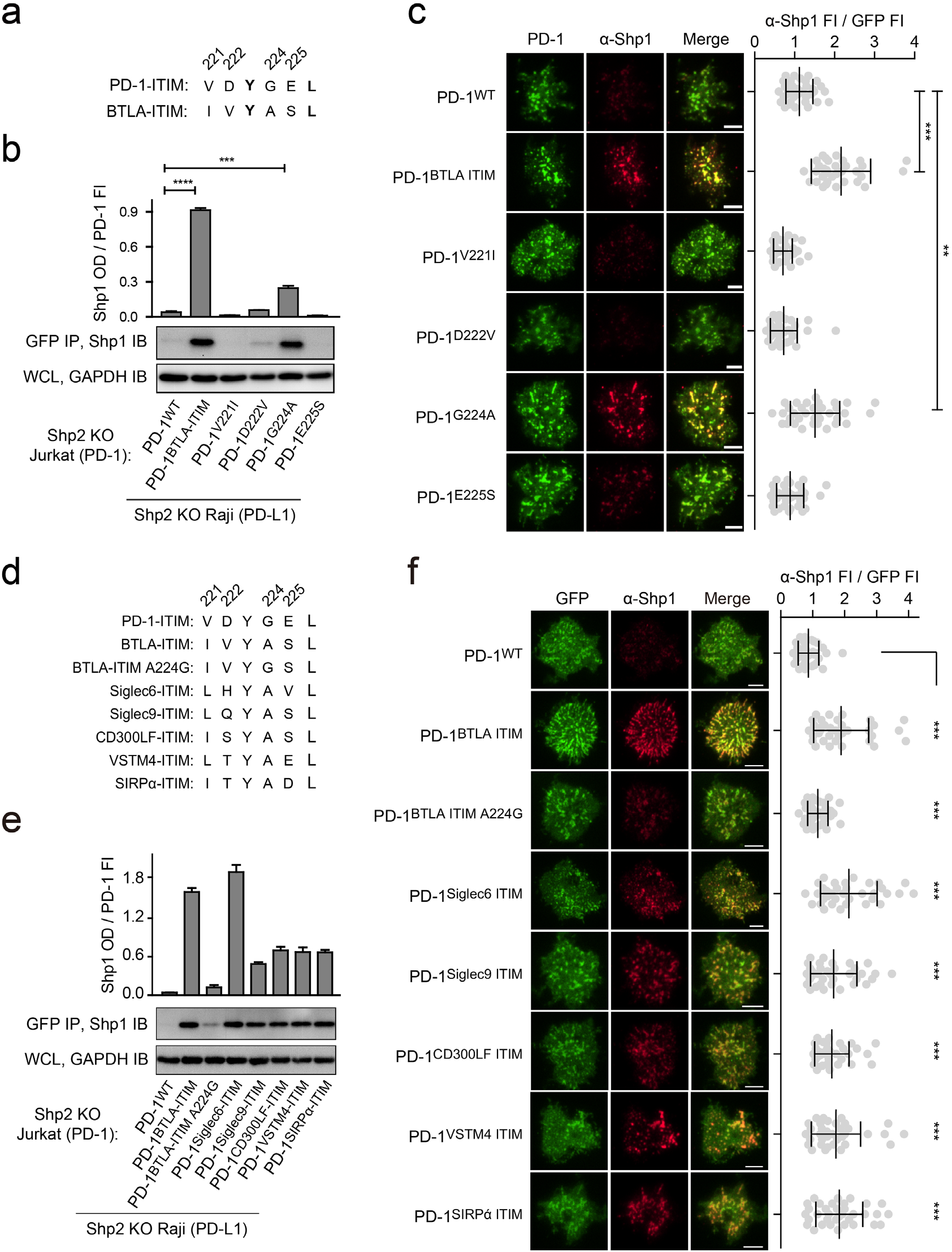
Glycine to alanine substitution at the pY+1 position of PD-1 ITIM promoted Shp1 recruitment. **a**, Cartoons depicting the amino acid (AA) alignment of PD-1-ITIM and BTLA-ITIM. **b**, Representative IBs showing the levels of Shp1 bound to mGFP-tagged PD-1 variants pulled down from the indicated co-culture lysates via GFP IP. GAPDH IB of WCL served as a loading control. Bar graphs summarize Shp1 OD normalized to the FI of each PD-1 variant, based on flow cytometry data in Extended Data Fig. 1d. Error bars are s.d. from three independent experiments. **c**, Left, representative TIRF images of both PD-1 (GFP) and endogenous Shp1 (stained with anti-Shp1) in an SLB-associated Shp2 KO Jurkat cell expressing indicated PD-1 variants. Right, dot plots summarizing anti-Shp1 FI normalized to GFP FI of 35 Jurkat cells under each condition (see Methods); Error bars: s.d.. Scale bars: 5 µm. **d**, Cartoons showing the AA sequences of ITIM of indicated receptors. **e**, Representative IBs showing the levels of Shp1 co-precipitated with mGFP-tagged PD-1 variants, with the original ITIM replaced by the indicated ITIM, from the indicated co-culture lysates via GFP IP. GAPDH IB of WCL served as a loading control. Bar graphs summarize Shp1 OD normalized to the FI of each PD-1 variant, based on flow cytometry data in Extended Data Fig. 1e. Error bars are s.d. from three independent experiments. **f**, Left, representative TIRF images of both PD-1 (GFP) and endogenous Shp1 (stained with anti-Shp1) in an SLB-associated Shp2 KO Jurkat cell expressing PD-1 variants with indicated ITIM. Right, dot plots summarizing anti-Shp1 FI normalized to GFP FI of 35 Jurkat cells under each condition (see Methods); Error bars: s.d.. Scale bars: 5 µm. **P < 0.01; ***P < 0.001; ****P < 0.0001; Student’s t-test

We next validated these findings using TIRF imaging of SLB-stimulated T cells. We observed PD-1 microclusters for all six PD-1 variants. Consistent with **Fig. 7c**, microclusters of PD-1^BTLA-ITIM^, but not PD-1^WT^ recruited Shp1 (**Fig. 8c**, rows 1 and 2). Among the single point mutants, only PD-1^G224A^ microclusters clearly recruited Shp1 (**Fig. 8c**, row 5), albeit less than did PD-1^BTLA-ITIM^ microclusters when Shp1 signal was normalized to the PD-1 signal (dot plot). In contrast, the other three single point mutants (PD-1^V221I^, PD-1^D222V^, and PD-1^E225S^) showed little to no Shp1 recruitment (**Fig. 8c**, rows 3, 4 and 6). These experiments suggested that the alanine residue at the pY+1 position of the ITIM motif promotes Shp1 binding.

Indeed, sequence alignment revealed that alanine is conserved at pY+1 position of ITIM in several inhibitory receptors (**Extended Data Fig. 5**), including Siglec-6, Siglec-9, CD300LF, VSTM4, and SIRPα, most of which reportedly recruit Shp1^26,38-41^. As expected, swapping the PD-1-ITIM by the ITIM of the foregoing inhibitory receptors (**Fig. 8d** and **Extended Data Fig. 1e**) significantly increased Shp1 recruitment to PD-1 immunoprecipitates and PD-1 microclusters, as compared to PD-1^WT^ (**Fig. 8e,f**).

### A medium-sized nonpolar residue at pY+1 position of the ITIM is required for Shp1 recruitment

The ability of somewhat conservative mutation (G224A) to induce PD-1:Shp1 binding was unexpected, however, given that alanine has a larger side chain than does glycine, we next sought to determine how the side chain property of pY+1 position influences PD-1:Shp1 interaction. Therefore, we mutated the glycine to a series of residues that differ in the size and polarity of their side chains. We established Jurkat cell lines that express each of the PD-1 mutants fused to a GFP tag, at comparable levels indicated by flow cytometry (**Extended Data Fig. 1f**). In the cell-SLB assay, we found that PD-1 mutants containing a medium-sized nonpolar residue at pY+1 position (G224A, G224V, G224L, G224I) most strongly recruited Shp1 (**Fig. 9a**, rows 3-6), a bulky or rigid nonpolar residue at this position (G224F, G224W, and G224P) decreased the Shp1 recruitment (**Fig. 9a**, rows 7-9). Indeed, plotting the Shp1 recruitment against the side chain volume of the five nonpolar residues, excluding the rigid proline, revealed a bell-shaped dependence that peaked at leucine and isoleucine (**Fig. 9b**). Finally, a polar or charged residue at this position failed to induce Shp1 recruitment, as observed for G224S, G224T, G224K and G224D mutants (**Fig. 9a**, rows 10-13). Taken together, these results revealed that a medium-sized nonpolar residue at pY+1 position of the ITIM is required for Shp1 recruitment. It is likely that Shp1-nSH2 contains a hydrophobic pocket that coordinates a medium-sized side chain of the pY+1 residue of ITIM.

**Fig. 9:**
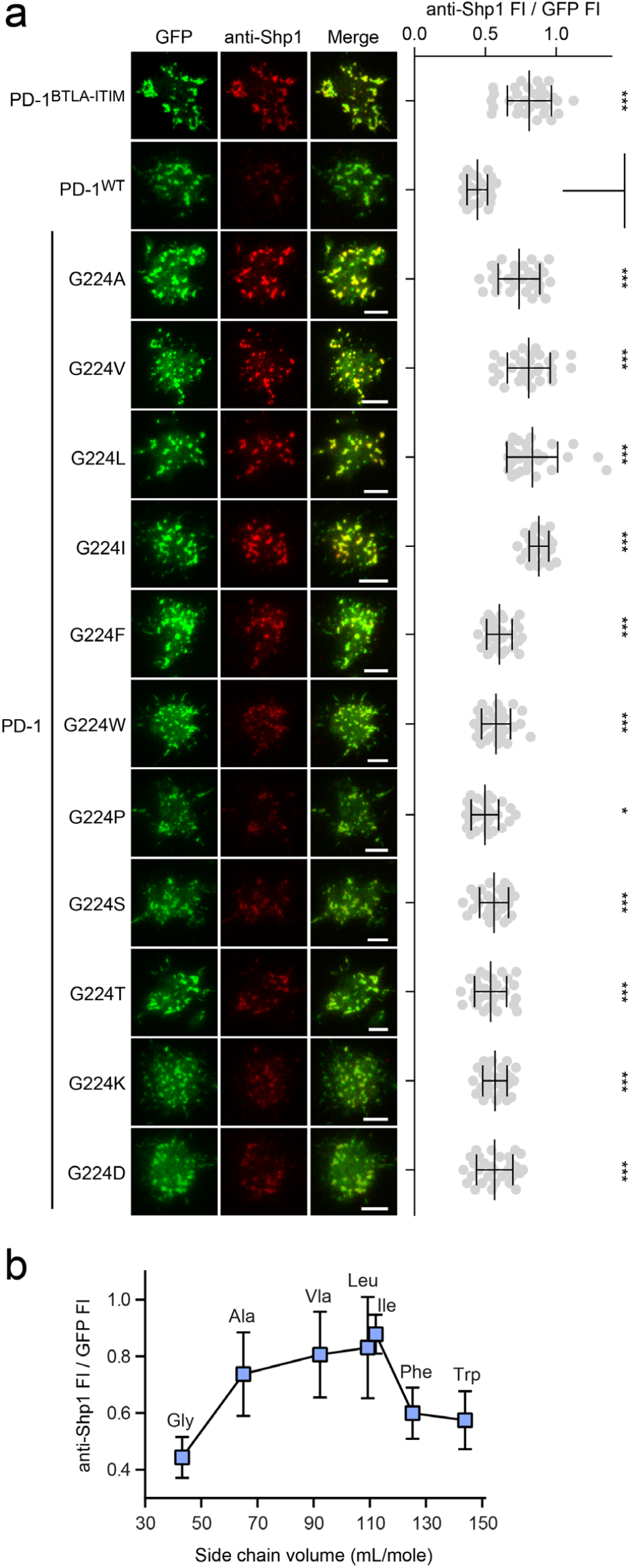
A medium-sized nonpolar residue at pY+1 position of the ITIM is optimal for Shp1 recruitment. **a**, Left, representative TIRF images of both PD-1 (GFP) and endogenous Shp1 (stained with anti-Shp1) in an SLB-associated Shp2 KO Jurkat cell expressing indicated PD-1 variants. Right, dot plots summarizing anti-Shp1 FI normalized to GFP FI of 30 Jurkat cells under each condition (see Methods); Error bars: s.d.. Scale bars: 5 µm. **b**, Normalized anti-Shp1 FI for a subset of PD-1 variants shown in A plotted against the nonpolar side chain volume of AA at the pY+1 position. Error bars: s.d., 30 cells. *P < 0.05; ***P < 0.001; ns, not significant; Student’s t-test.

## Discussion

Shp1 and Shp2 are key regulators of cell survival, proliferation, differentiation, and migration^29,42-45^. Co-expressed in hematopoietic cells, they operate as central effectors for immune inhibitory receptors that contain ITIM and ITSM. Dissecting the precise mechanism by which these receptors discriminate between Shp1 and Shp2 is required to predict and understand their “checkpoint” functions. In the present work, we combined biophysical, biochemical and cellular imaging approaches to investigate the specificity dichotomy of PD-1 and BTLA. Our data have revealed distinct properties between the SH2 domains of Shp1 and Shp2, and between the ITIMs of these two checkpoint receptors. We also report that the differential Shp1-binding activities of PD-1 and BTLA can be largely attributed to a single (pY+1) residue of their ITIMs: our data show that the polarity and size of this residue gate Shp1 recruitment in T cells.

In human genome, at least 32 receptors contain two or more ITIMs or ITSMs in tandem. Conceivably, the tandem pY motifs favor their bivalent interactions with tSH2 of Shp1 and Shp2. Mathematical modeling predicts that bivalent binding, even contributed by two weak bonds, can produce stable protein complexes due to the avidity effect and a reduction of off-rate of protein complexes^46^. In support of this notion, an earlier study showed that mutation of either ITIM or ITSM of BTLA abolishes its association with both Shp1 and Shp2^15^, a result that we have confirmed in the present study (**Fig. 5b**). The bivalent binding mode appears to be less strict in the case of PD-1, since its ITIM mutant retained the ability to co-IP Shp2^12,14,23,35,36^. A recent study suggested that Shp2 may crosslink two PD-1 molecules at the ITSM^36^, whereas other studies indicate that PD-1-ITIM contributes to Shp2 recruitment and activation^23,35,37^. In our hands, ITIM mutant recruited significantly less Shp2 than PD-1^WT^ in co-IP, microcluster enrichment and single molecule assays, supporting the involvement of ITIM, and the 1:1 bivalent mode as the dominant mode of PD-1:Shp2 interaction (**Fig. 2** and **Extended Data Fig. 2**). In both BTLA and PD-1, the ITIM tyrosine and ITSM tyrosine are separated by 25 residues^27^, and similar spacing are found in numerous ITIM/ITSM-containing receptors (**Supplementary Table 4**). This conserved distance may allow for simultaneous engagement of the ITIM and ITSM by the tSH2 of Shp1/Shp2 in the 1:1 bivalent mode. The bivalent binding likely occurs in a sequential fashion, in which the higher-affinity intermolecular contact precedes and converts the low affinity binding to a pseudo-intramolecular event, as supported by mathematical modeling^47^.

Our SPR data revealed certain promiscuities in the binding of their SH2 to ITIM and ITSM (**Figs. 3** and **6**). Theoretically, Shp1 and Shp2 may bind to PD-1 and BTLA in either a parallel or an antiparallel orientation. However, free energy calculations suggest parallel mode as the most stable form for both PD-1 and BTLA (**Supplementary Table 2**), consistent with a recent study on PD-1:Shp2 interaction^37^ and an earlier study on PECAM1:Shp2 interaction^48^. To our knowledge, all ITIM/ITSM receptors with the exception of SLAMF5 in human genome contain an ITIM N-terminal to ITSM (**Supplementary Table 4**). This spatial arrangement might allow these receptors to bind Shp1 and Shp2 in the parallel mode. The physiological significance of parallel binding is unclear, but it might increase the molecular reach of the PTPase domain^49,50^, or avoid potential steric clash with the plasma membrane.

Extensive genetic and biochemical evidence show that Shp1 and Shp2 differ in their physiological functions. Shp2 reportedly acts as both a positive and negative regulator whereas Shp1 is primarily known as a negative regulator in cell signaling^29,51^. The biochemical basis of their functional divergence is unknown. The most striking difference between Shp1 and Shp2, based on the current study, are their cSH2 domains: while Shp2-cSH2 binds to PD-1-ITSM with high affinity, Shp1-cSH2 exhibits weak to no binding to PD-1-ITSM (**Fig. 3a,b**). This distinction account for the undetectable PD-1:Shp1 association in T cells^23,24^, supported by our domain-swapping experiments (**Fig. 4**). Thus, PD-1 prefers Shp2 over Shp1 primarily based upon the stability of ITSM:cSH2 interaction. We speculate, in a more general sense, that the differing cSH2 domains of Shp1 and Shp2 might enable their recruitment to distinct signalosomes, leading to differing functional outcomes^29,51^. Indeed, Shp1-cSH2 and Shp2-cSH2 exhibit 51.9% homology in amino acid identities, and based on a recent nuclear magnetic resonance structure^37^, several residues implicated in PD-1-ITSM:Shp2-cSH2 interactions are altered in Shp1-cSH2.

Given the very weak affinity of the Shp1-cSH2 domain with the ITSM of both PD-1 and BTLA, we propose that receptors that stably recruit Shp1 must contain a strong docking site for Shp1-nSH2, such as the BTLA-ITIM (**Fig. 6a,b**). The PD-1-ITIM, however, might be too weak to support stable, bivalent PD-1:Shp1 binding. Moreover, because Shp1-nSH2 can bind intramolecularly to its catalytic domain to inhibit its PTPase activity^52,53^, a stronger ITIM might also allow the receptor to more efficiently release the auto-inhibition of Shp1.

Biochemical research on PD-1 has led to a consensus that ITSM is its primary docking site for Shp2^12,14,23,35,36^. Along this line, the current study shows that PD-1 uses ITSM to prefer Shp2 over Shp1 (**Fig. 3a,b**). Thus, ITSM is the ‘dominant hand’ of PD-1. In contrast, the ‘dominant hand’ of BTLA appears to be its ITIM. Our data shows that ITIM of BTLA serves as the primary docking site for Shp1, and allows BTLA to discriminate between Shp1 and Shp2 (**Fig. 6a,b**).

SH2 binding is contributed by both pY and its flanking residues^54,55^. Alanine is conserved at pY+1 position in the ITIMs of numerous immunoreceptors (**Extended Data Fig. 5**), suggesting its critical role in the physiological functions of these receptors. In this sense, PD-1 is an interesting exception: its ITIM appears the only ITIM that has a glycine at pY+1 position (**Extended Data Fig. 5**). Our study demonstrates that the pY+1 glycine inhibits Shp1 recruitment, and a single replacement of glycine to an alanine, valine, leucine or isoleucine was sufficient to induce PD-1:Shp1 association in T cells (**Figs. 8** and **9**). These results suggest that a medium-sized nonpolar residue at pY+1 position is a defining feature of a Shp1-docking site, specifically its nSH2 domain.

How could an alanine at pY+1 contribute to SH2 binding? In a reported crystal structure of NKG2A-ITIM:Shp1-cSH2 complex (PDB code: 2YU7), the methyl group of pY+1 alanine form hydrophobic contacts with nonpolar groups next to the pY-binding pocket within Shp1-cSH2. We speculate that similar methyl-mediated hydrophobic contacts likely stabilize BTLA-ITIM:Shp1-nSH2 interactions. A glycine at pY+1, as occurs in human PD-1-ITIM, likely prohibits such hydrophobic interactions due to the lack of a methyl group in its side chain. Leucine and isoleucine might further promote the interaction due to the more methyl groups in their side chains. Alternatively, the size of the pY+1 side chain might dictate the strength of the hydrophobic contact, a medium-sized residue (leucine, isoleucine, valine or alanine) might optimize the contact, whereas a small (glycine) or a bulky residue (phenylalanine or tryptophan) would inhibit the interaction due to a steric effect. Structural studies will be needed to definitively test these notions.

The result that leucine or isoleucine at pY+1 position enabled the most Shp1 recruitment (**Fig. 9**) suggests that the alanine-containing, naturally-occurring ITIMs are suboptimal for recruitment of Shp1, and modulating the side chain volume of pY+1 residue of ITIM may offer a means to tune Shp1 binding activities in cell therapies. However suboptimal binding affinities may also implicate an important regulatory role of the ITIM motif, for example a moderate level of Shp1/Shp2 recruitment may be utilized in some contexts to restrict tonic inhibitory signaling and promote cell fitness^56^. More generally, the diverse binding specificities of dual ITIM/ITSM motifs may enable more dynamic signaling pathways that are essential for homeostasis of the immune system.

## Acknowledgements

We thank P. Dennett and J. Zhang for critically reading the manuscript. T. Masubuchi is supported by the Human Frontier Science Program postdoctoral fellowship. This work was supported by R37 CA239072 from the National Institute of Health, a Searle Scholar Award from the Kinship Foundation, a Pew Biomedical Scholar Award from the Pew Charitable Trusts to E. Hui.

## Author contributions

X.X., T.M., and E.H. designed the project. X.X., T.M. and Y.Z. generated plasmids, proteins and cell lines. X.X. conducted the co-IP experiments. T.M. conducted the cell-SLB assays, SPR experiments, and single molecule imaging. X.X., T.M., and E.H. wrote the manuscript.

## Methods

### Reagents

Biotin anti-human CD3ε (clone OKT3, BioLegend, #317320), human PD-L1-His (Sino Biological, #10084-H08H), human ICAM-1-His (Sino Biological, #10346-H08H), and streptavidin (Invitrogen, #S888) were used to functionalize SLBs. Anti-Shp1 (Life Technologies, #3H20L13), anti-Shp2 (BD Biosciences, #101720), and F (ab’) 2-Goat anti-Rabbit IgG (H+L) secondary antibody, alexa fluor 568 (Invitrogen, #A21069) were used in the immunostaining for the cell-SLB assays. Anti-Phosphotyrosine (clone pY20, Sigma-Aldrich, #P4110-1MG), anti-GFP (Invitrogen, #A-6455), anti-Shp1 (Proteintech, #24546-1-AP), anti-Shp2 (Santa Cruz Biotechnology, #sc-7384), and Anti-GAPDH (Proteintech, #10494-1-AP) were used in western blots. Phycoerythrin (PE) anti-human BTLA (clone MIH26, BioLegend, #344505) and Pacific Blue anti-human PD-1 (clone EH12.2H7, BioLegend, #329916), were used in the staining of Jurkat or Raji cells for flow cytometry.

### Cell cultures

HEK293T cells were maintained in DMEM medium (Genesee Scientific, #25-501) supplemented with 10% fetal bovine serum (Omega Scientific, #FB-02) and 1% 100 × Penicillin-Streptomycin (GE Healthcare, #SV30010) at 37 °C / 5% CO_2_. Jurkat and Raji cells were maintained in RPMI-1640 medium (Corning, #10-041-CM) supplemented with 10% fetal bovine serum, 100 U/mL of Penicillin, and 100 µg/mL of Streptomycin) at 37 °C / 5% CO_2_.

### Recombinant proteins

For SPR assays in **Figs. 3a** and **6a**, human BTLA^ICD^ (aa 190-289), human PD-1^ICD^ (aa 194-288), and their tyrosine mutants (BTLA^FFYF^, BTLA^FFFY^, PD-1^FY^, PD-1^YF^) were expressed with an N-terminal His_10_ tag in *Escherichia coli* using the pET28A vector, and purified using Ni-NTA agarose (ThermoFisher, #88223) as described^24^. His_10_ tagged human protein tyrosine kinase Fyn was expressed in the Bac-to-Bac baculovirus system and purified using Ni-NTA agarose. SNAP-Shp1-nSH2, SNAP-Shp1-nSH2, SNAP-Shp2-nSH2, and SNAP-Shp2-cSH2 were expressed with an N-terminal GST tag followed by a PreScission recognition sequence (LEVLFQGP), in *Escherichia coli* via the pGEX6p-2 vector. For the single molecule imaging assay in **Extended Data Fig. 2**, all proteins were expressed with an N-terminal GST tag followed by a PreScission recognition sequence (LEVLFQGP), in *Escherichia coli* via the pGEX6p-2 vector. These included SNAP-Shp1-tSH2, SNAP-Shp2-tSH2, as well as the ICD of PD-1 WT or its tyrosine mutants fused with an N terminal Avi-tag (GLNDIFEAQKIEWHE) and a C-terminal SNAPf tag (Avi-PD-1^YY^-SNAPf, Avi-PD-1^FY^-SNAPf, Avi-PD-1^YF^-SNAPf). All GST fusion proteins were purified using Glutathione Agarose 4B (Gold Biotechnology, #G-250-50), and eluted with HEPES buffered saline (HBS, 50 mM HEPES, 150 mM NaCl, 0.5 mM TCEP (Gold Biotechnology, #TCEP10), pH 7.5) containing 20 units/mL 3C protease to remove the GST tag. After elution, SNAP-Shp1-tSH2, SNAP-Shp2-tSH2 were further labeled with SNAP ligand-JF646 (Janelia Research Campus (HHMI), #Janelia 2014-013) at 4 °C overnight. Avi-PD-1^YY^-SNAPf, Avi-PD-1^FY^-SNAPf, Avi-PD-1^YF^-SNAPf were further labeled with SNAP ligand-JF549 (Janelia Research Campus (HHMI), #Janelia 2014-013) and biotin in the presence of 1 mM Biotin (Sigma-Aldrich, #B4501), 1 μM BirA and 10 mM ATP (Gold Biotech, #A-081-100) at 4 °C overnight. All affinity-purified proteins were subjected to gel filtration chromatography using HBS containing 10% glycerol and 1 mM TCEP. The monomer fractions were pooled, snap frozen and stored at - 80 °C in small aliquots. Gel filtration standards (Bio-Rad, #1511901) were run to confirm the sizes of eluted proteins.

### Phosphorylation of recombinant PD-1 and BTLA ICD proteins

Purified His_10_-PD-1, His_10_-BTLA and Avi-PD-1-SNAPf proteins were incubated with 50 nM purified Fyn, 2 mM ATP, and 10 mM Na_3_VO_4_ at RT for 6 hours to achieve full phosphorylation, as indicated by the complete shift of electrophoretic mobility on sodium dodecyl sulfate polyacrylamide gel electrophoresis (SDS-PAGE). Monomeric form of these pre-phosphorylated proteins were purified using gel filtration chromatography in HBS containing 10% glycerol and 1 mM TCEP, and stored in - 80 °C in aliquots.

### Shp2 antibody labeling

For fluorescent staining of Shp2 in T cells, anti-Shp2 was fluorescently labeled using Alexa Fluor 647 NHS ester (ThermoFisher, #A37573), and unreacted chemicals were removed using Zeba Spin Desalting Columns (ThermoFisher, #89890) following manufacturer’s instructions.

### Cell lines

Jurkat E6.1 cells were provided by Dr. Arthur Weiss (University of California San Francisco). HEK293T cells and Raji cells were provided by Dr. Ronald Vale (University of California San Francisco). Shp2 KO Jurkat, Shp1/Shp2 DKO Jurkat, Raji (PD-L1-mCherry), Raji (HVEM-mRuby2), Shp2 KO Raji (PD-L1-mCherry), and Shp2 KO Raji (HVEM-mRuby2) cells were generated in our previous study^24^. Each gene of interest was introduced into Jurkat cells via lentiviral transduction, as described previously^24^. Briefly, each cDNA was cloned into a pHR vector backbone, and co-transfected with pMD2.G and psPAX2 packaging plasmids into HEK293T cells using polyethylenimine (PEI, Fisher Scientific, #NC1014320). Virus-containing supernatants were harvested at 60-72 h post-transfection. WT Jurkat cells, Shp2 KO Jurkat cells and Shp1/Shp2 DKO Jurkat cells were resuspended with the desired virus supernatant, centrifuged at 35 °C, 1000 × g for 30 min, and incubated overnight at 37 °C, 5% CO2 before replacing the virus supernatant with complete RPMI-1640 medium.

### Jurkat-Raji co-culture assay and immunoprecipitation

For **Figs. 1, 2a, 4c, 5b, 7b, 8b, 8e, Extended Data Figs. 3b** and **4b**, Jurkat cells were starved in serum free RPMI medium at 37 °C for 3 h prior to co-culture. Raji cells were pre-incubated with 30 ng/mL SEE (Toxin Technologies, #ET404) in RPMI medium for 30 min at 37 °C. In order to avoid Shp2 competition from Raji cells, Shp2 KO Raji cells were used in **Figs. 4c, 7b, 8b, 8e** and **Extended Data Fig. 4b**. Afterwards, four million SEE-loaded Raji cells and four million Jurkat cells were precooled on ice and mixed in a 96-well plate. After centrifugation at 300 × g for 1 min at 4 °C to initiate Raji-Jurkat contact, cells were immediately transferred to a 37 °C water bath. At 5 min or the time points indicated in the figures, Raji-Jurkat conjugates were lysed with HBS containing 5% glycerol, detergent (1% NP-40), protease inhibitor (1 mM PMSF), and phosphatase inhibitors (10 mM Na_3_VO_4_ and 10 mM NaF). GFP-tagged PD-1 variants or BTLA variants were IP’ed from the lysate using GFP-Trap (Chromotek, #gta-20). Equal fractions of the IP samples were subjected to SDS-PAGE and blotted with indicated antibodies.

### Flow cytometry

Flow cytometry was conducted in an LSRFortessa cell analyzer (BD Biosciences). For **Fig. 5a, Extended Data Figs. 1, 3a** and **4a**, indicated Jurkat cells were washed with PBS and analyzed after staining with PE anti-human BTLA (MIH26) or Pacific Blue anti-human PD-1 (EH12.2H7). For **Extended Data Fig. 1b**, mCherry levels were measured using a FACSAria cell sorter (BD Biosciences) due to a lack of 561 nm laser in the LSRFortessa cell analyzer. Data analyzed using FlowJo (FlowJo, LLC).

### SLB preparation and functionalization

SLBs were formed on glass-bottomed 96-well plate (DOT Scientific Inc, #MGB096-1-2-LG-L). Briefly, the plate was cleaned with 2.5 % Hellmanex (Sigma-Aldrich, #Z805939-1EA), etched with 5N NaOH, and used for SLB formation as previously described ^57^. Briefly, small unilamellar vesicles (SUVs) derived from dried lipid film containing 95.5% POPC (Avanti Polar Lipids, #850457C), 2% biotin-DPPE (Avanti Polar Lipids, #870285P), 2% DGS-NTA-Ni (Avanti Polar Lipids, #790404C) and 0.1% PEG 5000-PE (Avanti Polar Lipids, #880230C) were added onto freshly treated plates to form SLBs. The SLBs were rinsed with wash buffer (1x PBS containing 0.1% BSA) and mixed with 1 μg/mL streptavidin, 0.1 nM His-tagged human PD-L1 ectodomain, and 3 nM His-tagged human ICAM-1 ectodomain at 37 °C for 1 h. Afterward, the SLBs were rinsed with wash buffer and further incubated with 5 μg/mL biotin anti-human-CD3ε at 37 °C for 30 min, followed by three rinses with wash buffer and three rinses with imaging buffer (20 mM HEPES pH 7.5, 137 mM NaCl, 5 mM KCl, 1 mM CaCl_2_, 2 mM MgCl_2_, 0.7 mM Na_2_HPO_4_, 6 mM D-Glucose).

### Cell-SLB image acquisition and analysis

Jurkat cells were resuspended in imaging buffer and overlaid onto freshly formed PD-L1/ICAM/Okt3-functionalized SLBs. After 5 min incubation at 37 °C, SLB-bound cells were overlaid with 2% paraformaldehyde (PFA, Fisher Scientific, #50980494), and incubated at room temperature (RT) for 15 min for fixation. SLB-associated, PFA-treated cells were washed with blocking buffer (1x PBS containing 3% BSA), and permeabilized with 1x PBS containing 3% BSA and 0.1% Saponin at RT for 30 min. To observe mCherry-tagged Shp1^WT^ or domain swapping mutants (**Fig. 4b**), cells were directly imaged at both GFP (488 nm) and mCherry (561 nm) channels. To observe endogenous Shp2 (**Fig. 2b**), the permeabilized cells were stained with Alexa-Fluor-647-labeled anti-Shp2 at 4 °C for 16 hours, followed by fixation with 4% PFA. To observe endogenous Shp1 (**Figs. 7c, 8c, 8f**, and **9a**), the permeabilized cells were stained with anti-Shp1 at 4 °C for 16 hours, followed by fixation with 4% PFA, and further staining with Alexa-Fluor-568-labeled anti-rabbit IgG at RT for 1 hour and another treatment with 4% PFA. The fluorescent cell images were acquired on a Nikon Eclipse Ti TIRF microscope equipped with a 100x Apo TIRP 1.49 NA objective lens, controlled by the Micro-Manager software^58^. Fiji^59^ was used to quantify the degree of recruitment of Shp1 and Shp2 to PD-1 microclusters. Mask images identifying the area of PD-1 microclusters were generated by applying the “subtract background” command to PD-1 (mGFP) images using the default setting. The fluorescent signals of anti-Shp1, anti-Shp2 and PD-1 (GFP) in the masked overlaid images were measured, and used to calculate the anti-Shp1 FI/GFP FI ratio and Shp2/PD-1 ratio for each cell.

### Surface plasmon resonance (SPR) assay

For **Figs. 3a** and **6a**, direct interaction between individual SH2 (nSH2 or cSH2) of Shp1 or Shp2 and BTLA ICD, PD-1 ICD, or their tyrosine mutants (BTLA^FFYF^, BTLA^FFFY^, PD-1^FY^, PD-1^YF^) was monitored by OpenSPR (Nicoya) equipped with Ni sensor chip (Nicoya, #SEN-AU-100-10-NTA). Pre-phosphorylated His_10_-tagged PD-1 ICD or BTLA ICD was immobilized onto a Ni sensor chip to achieve approximately 1500 RU by following the Ni sensor wizard in OpenSPR software. SH2 of interest was diluted in running buffer (20 mM HEPES, 150 mM NaCl, 5 mM Imidazole (Sigma-Aldrich, #I202), 0.05 % Tween-20, 10 % Glycerol, pH 7.5), and injected. The association and dissociation phases of SH2 were monitored at flow rate of 20 μL/min. The Ni sensor chip was regenerated with 50 mM NaOH before injecting the next SH2. Sensorgrams were analyzed using the “Evaluate EC50” method in TraceDrawer software (Ridgeview Instruments).

### Single molecule imaging assay

For **Extended Data Fig. 2**, surface-passivated coverslips and slide glasses used in single molecule imaging assays were prepared and assembled as previously described^60^. Surface-passivated glass chambers were incubated with blocking buffer (50 mM HEPES, 150 mM NaCl, 0.1 % BSA) at RT for 5 min and with 0.5 μM Streptavidin in blocking buffer at RT for 5 min, followed by two washes with blocking buffer. The glass chambers were further incubated with 10 pM pre-phosphorylated, biotinylated, JF549-labeled PD-1^YY^, PD-1^FY^, or PD-1^YF^, and incubate at RT for 5 min. The unbound proteins were removed with blocking buffer and JF646-labeled SH2 proteins were injected to the chambers. The fluorescent images were acquired at 20 Hz on a Nikon Eclipse Ti TIRF microscope equipped with a 100x Apo TIRP 1.49 NA objective lens, controlled by the Micro-Manager software^58^.

### Fluorescent intensity distribution analysis and photobleaching analysis

For **Extended Data Fig. 2c,d**, JF646-labeled PD-1 attached on coverslips were illuminated with the 50 mW 488 nm laser for photobleaching observation. The initial positions of fluorescent spots for JF646-labeled PD-1 were determined by the “ThunderSTORM” plugin ^61^ in Fiji. The fluorescent intensities at each determined position in initial images were measured and plotted into histogram fit with Gaussian distributions, and those of whole image stacks were measured for fluorescent trajectories.

### Fluorescent trajectory analysis and colocalization analysis

For **Extended Data Fig. 2d,f**, the positions of fluorescent spots for JF549-labeled PD-1 were determined by the “ThunderSTORM” plugin in Fiji. Likewise, the fluorescent movies for JF646-labeled tSH2 proteins were Z-projected to “maximum intensity” and analyzed by the “ThunderSTORM” plugin to determine the positions where tSH2 proteins bound. The positions detected in both JF549 and JF646 channels were determined as spots representing PD-1 molecules that recruited tSH2 protein at any given time during the image acquisition. The fluorescent intensities of JF646 within those areas were measured to fluorescent trajectories of tSH2 proteins. The bound and unbound states of tSH2 proteins in the fluorescent trajectories were estimated with Hidden Markov Model in a custom written Python script. The colocalization rates of PD-1 and tSH2 protein were calculated by dividing the total number of PD-1 spots by the number of PD-1 spots that recruited SH2 proteins within 0.5 seconds of image acquisitions.

### Quantification and statistical analysis

Data were shown as mean ± s.d., and number of replicates were indicated in figure legends. Curve fitting and normalization were performed in GraphPad Prism 8 (GraphPad). Statistical significance was evaluated by Student’s t-test (*, p < 0.05; **, p < 0.01; ***, p < 0.001; ****, p < 0.0001; ns, not significant). Data with p < 0.05 are considered statistically significant.

## Supplementary information for

**Extended Data Fig. 1.**
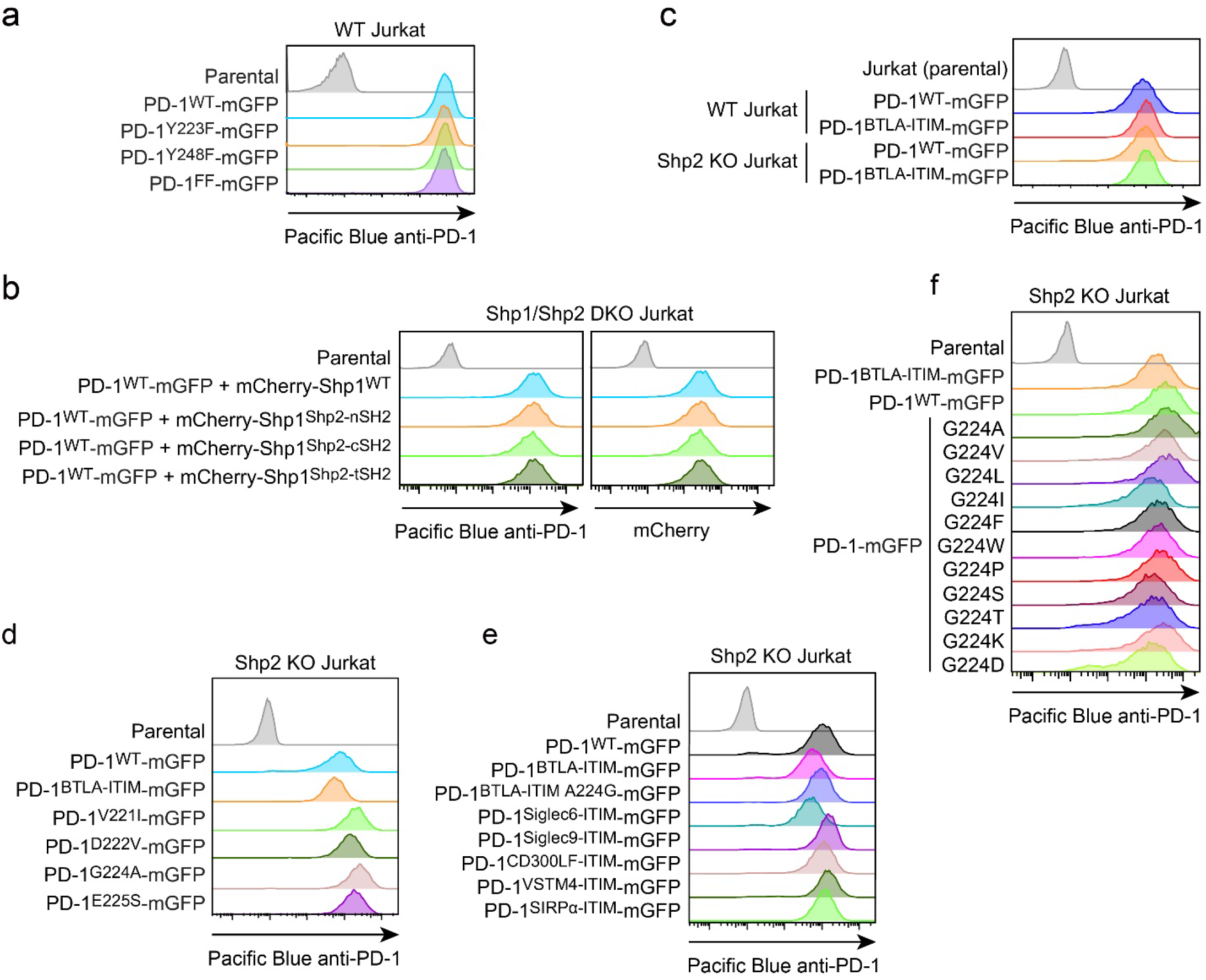
Flow cytometry histograms for Figs. 2, 4, and 7-9. **a**, Flow cytometry histograms showing cell surface expressions of PD-1 variants in indicated Jurkat cells. Related to Fig. 2. **b**, Flow cytometry histograms showing cell surface expression of PD-1WT and total expressions of indicated Shp1 variants in Shp1/Shp2 DKO Jurkat cells. Related to Fig. 4. **c**, Flow cytometry histograms showing cell surface expression of indicated PD-1 variants in WT or Shp2 KO Jurkat cells. Related to Fig. 7. **d**-**f**, Flow cytometry histograms showing cell surface expression of indicated PD-1 variants in Shp2 KO Jurkat cells. Related to Fig. 8a-c, Fig. 8d-f and Fig. 9, respectively.

**Extended Data Fig. 2.**
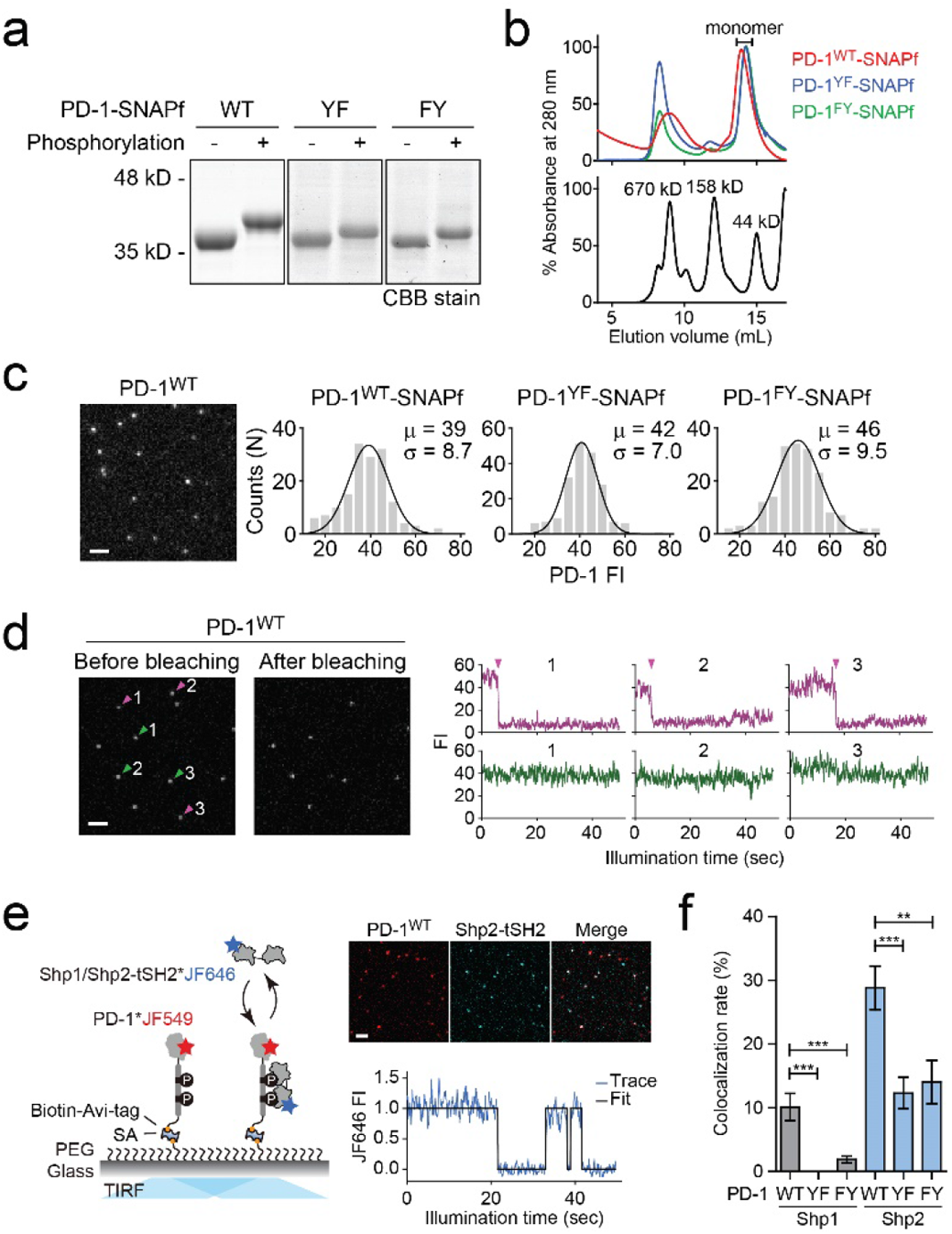
Single molecule imaging monitoring Shp1/Shp2-tSH2 binding to PD-1 variants. a, Coomassie-stained SDS-PAGE images showing the shift of electrophoretic mobilities of indicated PD-1 variants after phosphorylation. **b**, Size exclusion chromatograms of phosphorylated PD-1 variants (top) and protein standards with their molecular weights labeled in kilo Delton (kD, bottom). The monomeric fractions used for single molecule imaging assays are indicated. **c**, Left, a representative TIRF image showing JF646-labeled PD-1 single molecules. Right, histograms summarizing the FI of individual molecules of PD-1 in the TIRF image. Mean (µ) and s.d. (σ) of the Gaussian fit are indicated distribution. Scale bar: 2 µm. **d**, Left, representative TIRF images showing JF646-labeled PD-1 single molecules before and after photobleaching. The bleached and un-bleached spots were indicated with purple and green arrowheads, respectively. Right, fluorescence time trajectories of bleached (purple) and non-bleached (green) spots, with the bleaching moments indicated with arrow heads. Scale bar: 2 µm. **e**, Left, a cartoon depicting the setup of single molecule imaging assay. Each of the pre-phosphorylated, biotinylated, and JF549-labeled PD-1 variants was immobilized onto a biotin-PEG coated coverslip. The FIs of PD-1 molecules and the bound JF646-labeled Shp1/Shp2-tSH2 were monitored by TIRF-M. Right, representative TIRF images showing the PD-1^WT^ and Shp2-tSH2 single molecules (top), and a representative time trajectory showing Shp2-tSH2 binding and unbinding from PD-1^WT^ (bottom). Scale bar: 2 µm. **f**, Bar graphs summarizing the % of PD-1 molecules associated with Shp1-tSH2 or Shp2-tSH2 (see Methods). Error bars are s.d. from three independent experiments. **P < 0.01; ***P < 0.001; Student’s t-test.

**Extended Data Fig. 3.**
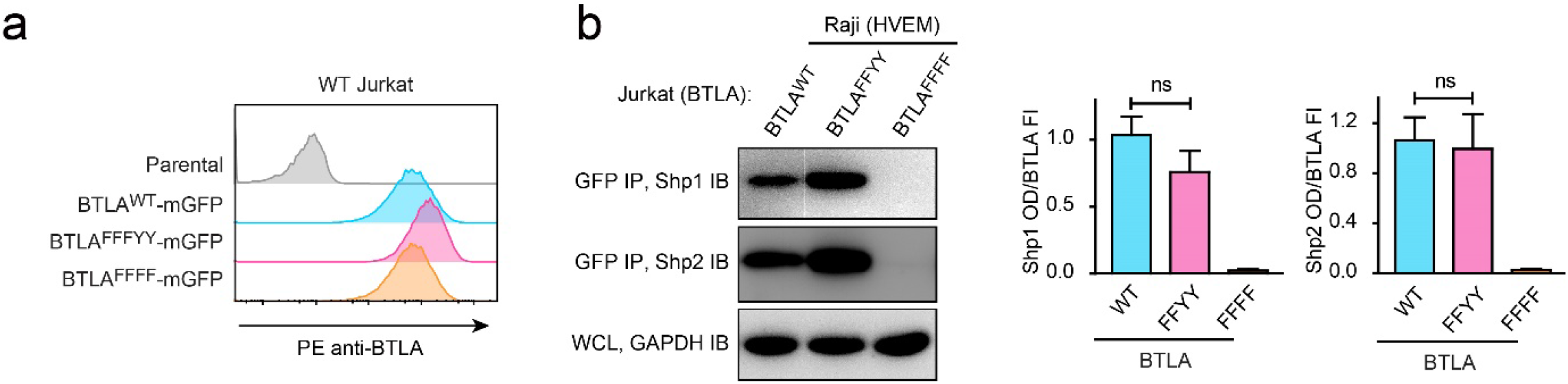
Y226 and Y243 are dispensable for Shp1/Shp2 recruitment by BTLA. **a**, Flow cytometry histograms showing BTLA surface expressions in the indicated Jurkat cells. **b**, Representative IBs showing the levels of Shp1 and Shp2 bound to the mGFP-tagged BTLA variants captured by GFP IP. GAPDH IB of the WCL served as a loading control. Right, bar graphs summarizing Shp1 OD and Shp2 OD normalized to the FI of the corresponding BTLA variants, based on flow cytometry data in A. Error bars are s.d. of three independent experiments. ns, not significant in Student’s t-test.

**Extended Data Fig. 4.**
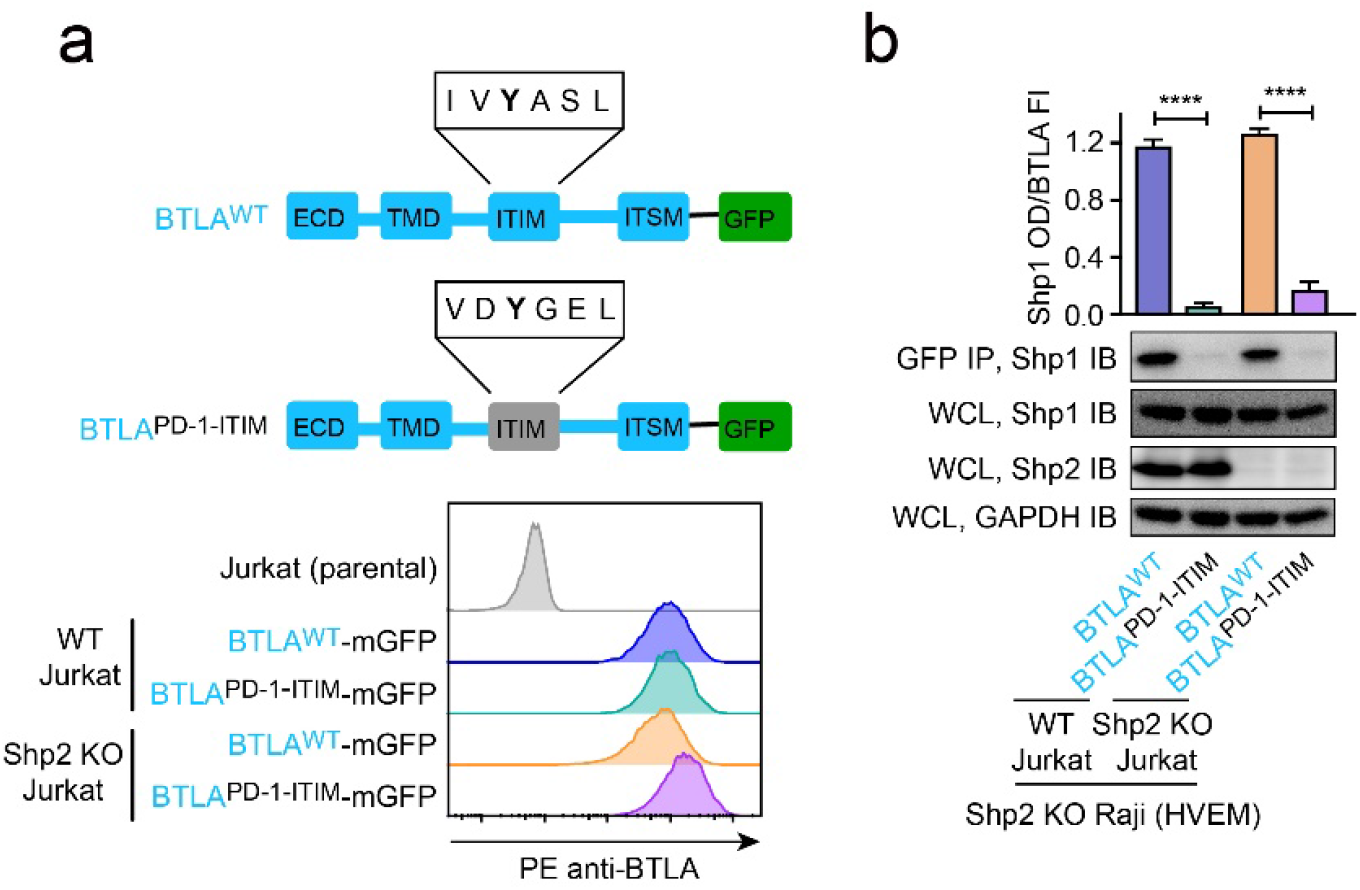
Replacing BTLA-ITIM with PD-1-ITIM abolished BTLA:Shp1 interaction in Jurkat cells. **a**, Upper, cartoons depicting the domains and motifs of BTLA^WT^-mGFP and BTLA^PD-1 ITIM^-mGFP. Lower, flow cytometry histograms showing the cell surface expressions of BTLA variants in the indicated Jurkat cells. **b**, Representative IBs showing the levels of Shp1 bound to mGFP-tagged BTLA variants pulled down from the indicated co-culture lysates via GFP IP. Shp1 IB and Shp2 IB of the WCL indicate their input. GAPDH IB of WCL serves as a loading control. The bar graphs on top summarizes Shp1 OD normalized to the FI of each BTLA variant, based on flow cytometry data in A. Error bars are s.d. from three independent experiments. ****P < 0.0001; Student’s t-test.

**Extended Data Fig. 5.**
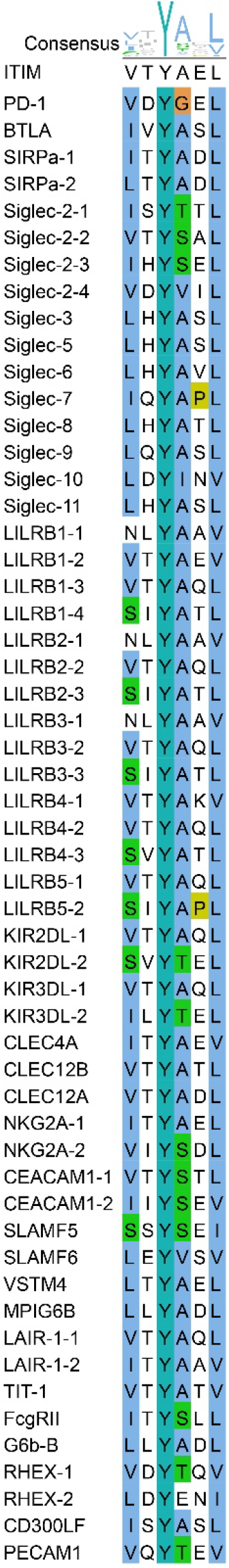
Sequence alignment of ITIMs in immune receptors. ITIM sequences in indicated immune receptors were aligned and visualized using ClustalW and JalView. Consensus amino acids at each position in ITIM are indicated at the top.

**Supplementary Table 1.** *K*_d_ values of individual SH2 of Shp1/Shp2 and phosphorylated ITIM/ITSM of PD-1/BTLA; mean±s.d. (n=3).

| $K_d$ (μM) | PD-1-ITIM | PD-1-ITSM | BTLA-ITIM | BTLA-ITSM |
| --- | --- | --- | --- | --- |
| Shp1-nSH2 | 0.27 ± 0.1 | 0.083 ± 0.020 | 0.064 ± 0.012 | n.d. |
| Shp1-cSH2 | n.d. | 1.7 ± 1.4 | 1.8 ± 0.20 | 0.86 ± 0.31 |
| Shp2-nSH2 | 0.38 ± 0.10 | 0.14 ± 0.049 | 0.34 ± 0.082 | 2.1 ± 1.1 |
| Shp2-cSH2 | 1.4 ± 0.57 | 0.10 ± 0.029 | 1.1 ± 0.54 | 1.1 ± 0.19 |

n.d.: not detected

**Supplementary Table 2.** ΔG of PD-1/BTLA interactions with Shp1/Shp2-tSH2 in a parallel or an anti-parallel mode.

| $\Delta G$ (kJ)<sup>a</sup> | Parallel | Anti-Parallel |
| --- | --- | --- |
| PD-1:Shp1tSH2 | -70.4 | -40.4 |
| PD-1:Shp2tSH2 | -76.6 | -72.5 |
| BTLA:Shp1tSH2 | -75.7 | -32.8 |
| BTLA:Shp2tSH2 | -70.9 | -66.4 |

a: $\Delta G = RT \cdot \ln(K_d)$
T=298.15 K, R is molar gas constant, $R=8.314 \times 10^{-3}$ kJ/K/M

**Supplementary Table 3.**
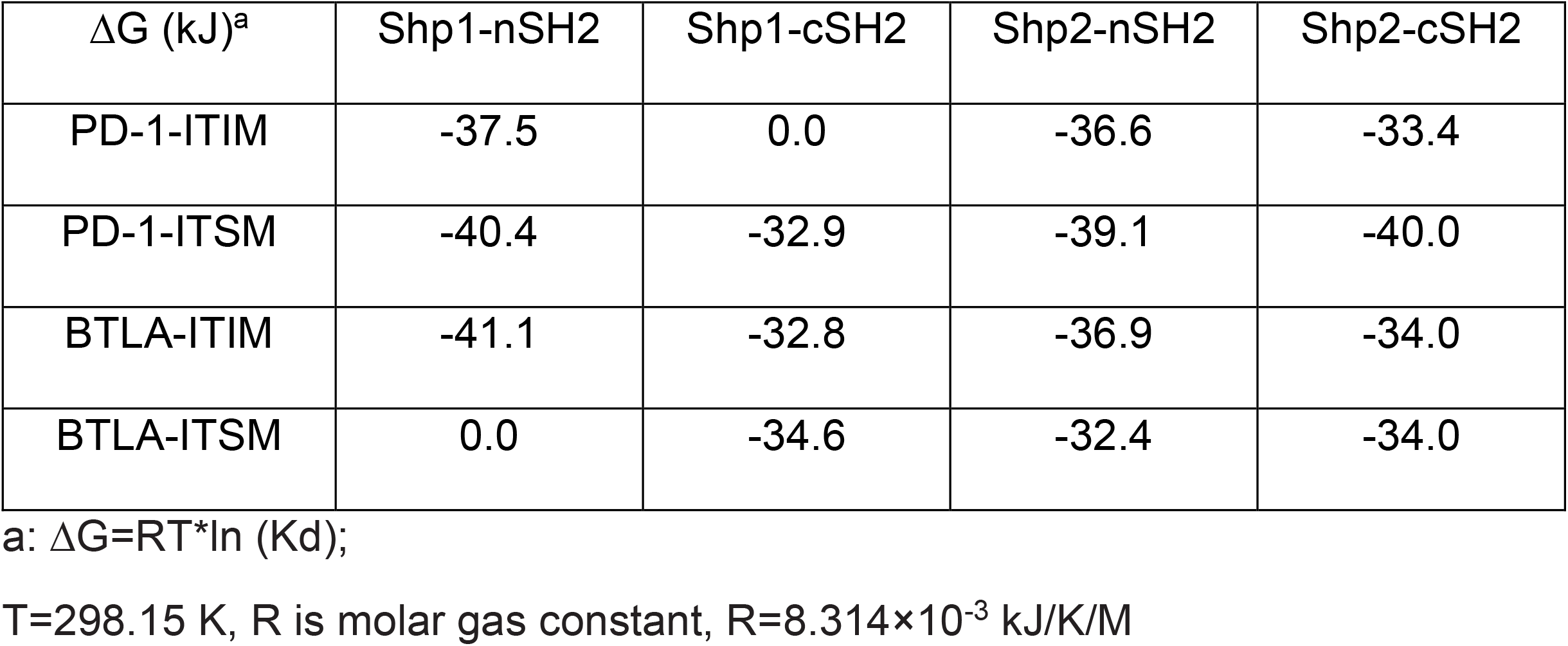
ΔG of ITIM/ITSM and SH2 bindings.

| $\Delta G$ (kJ)<sup>a</sup> | Shp1-nSH2 | Shp1-cSH2 | Shp2-nSH2 | Shp2-cSH2 |
| --- | --- | --- | --- | --- |
| PD-1-ITIM | -37.5 | 0.0 | -36.6 | -33.4 |
| PD-1-ITSM | -40.4 | -32.9 | -39.1 | -40.0 |
| BTLA-ITIM | -41.1 | -32.8 | -36.9 | -34.0 |
| BTLA-ITSM | 0.0 | -34.6 | -32.4 | -34.0 |

a: $\Delta G = RT \cdot \ln(K_d)$ ;
T=298.15 K, R is molar gas constant, $R=8.314 \times 10^{-3}$ kJ/K/M

**Supplementary Table 4.** Immune receptors harboring both ITIM and ITSM.

| Receptor | ITIM Y position<br>(AA) | ITSM Y position<br>(AA) | ITIM-ITSM distance (AA)<sup>a</sup> |
| --- | --- | --- | --- |
| PD-1 | 223 | 248 | 25 |
| BTLA | 257 | 282 | 25 |
| Siglec-3 | 340 | 358 | 18 |
| Siglec-5 | 520 | 544 | 24 |
| Siglec-6 | 426 | 446 | 20 |
| Siglec-9 | 433 | 456 | 23 |
| Siglec-11 | 644 | 668 | 24 |
| CD300LF | 249 | 284 | 35 |
| VSTM4 | 289 | 310 | 21 |
| SIIRPα | 429 | 453 | 24 |
|  | 470 |  | -16 |
|  | 496 |  | -43 |
| SLAMF5 | 341 | 279 | -62 |
|  |  | 316 | -25 |
| SLAMF6 | 274 | 285 | 11 |
|  |  | 309 | 35 |
| PECAM1 | 690 | 713 | 23 |
| MPIG6B | 211 | 237 | 26 |

AA: amino acids
ITIM: immunoreceptor tyrosine-based inhibition motif
ITSM: immunoreceptor tyrosine-based switch motif
a: Calculated as (ITSM Y position)-(ITIM Y position)

